# Reciprocal interactions between the gut microbiome and mammary tissue mast cells promote metastatic dissemination of HR^+^ breast tumors

**DOI:** 10.1101/2021.12.23.474065

**Authors:** Tzu-Yu Feng, Francesca N. Azar, Claire Buchta Rosean, Mitchell T. McGinty, Audrey M. Putelo, Sree Koli, Natascia Marino, Rana German, Ram Podicheti, Sally A. Dreger, Wesley J. Fowler, Stephanie Greenfield, Stephen D. Robinson, Melanie R. Rutkowski

## Abstract

Establishing commensal dysbiosis, defined as an inflammatory gut microbiome with low biodiversity, prior to breast tumor initiation, enhances early dissemination of hormone-receptor positive (HR^+^) mammary tumor cells. Here, we sought to define mammary tissue mediators of dysbiosis-induced tumor dissemination. We found that commensal dysbiosis increased both the frequency and profibrogenicity of mast cells in the mammary tissue, a phenotypic change that persisted after tumor implantation. Fibroblast activation and tissue remodeling associate with enhanced breast tumor metastasis. We employed pharmacological and adoptive transfer approaches to demonstrate that mammary tissue mast cells from dysbiotic animals enhances dissemination of HR^+^ tumor cells. Collagen levels in mammary tissues from HR^+^ breast cancer patients correlated with mast cell abundance, suggesting clinical relevance of mast cell-mediated fibroblast activation. Together, these data demonstrate that a gut-mast cell axis exists that induces fibroblast activation and orchestrates early dissemination of HR^+^ breast tumors.

**Significance:** Our study defines the mechanism by which an inflammatory gut microbiome facilitates HR^+^ breast tumor cell dissemination. We establish that gut commensal dysbiosis triggers mammary tissue mast cells to facilitate early metastatic dissemination. These findings highlight a novel gut microbiome-innate immune cell axis involved in negative breast cancer outcomes.

## Introduction

Despite successful first-line treatments, a substantial population of hormone-receptor positive (HR^+^: estrogen- and/or progesterone-receptor^+^, HER2^−^) breast cancer patients develops metastasis and succumb to their disease. According to the Surveillance, Epidemiology and End Results (SEER) database, HR^+^ breast cancer is the most common molecular subtype (73.1%) of the metastatic breast cancer population in the US^1^. Well-established treatment strategies have significantly increased long-term survival of women diagnosed with HR^+^ breast cancer, but accurate prediction of patient recurrence remains a significant barrier to reducing the mortality associated with metastatic breast cancer.

Using a novel murine model of HR^+^ breast cancer^2–4^, we recently demonstrated that gut commensal dysbiosis (an inflammatory gut microbiome with low biodiversity) established prior to breast tumor initiation enhances metastatic dissemination of HR^+^ tumors while primary tumor growth remained unchanged^5^. This was one of the first studies to demonstrate that the gut microbiome acts as a host-intrinsic mediator of breast tumor metastasis. Sequencing of the gut microbiome from patients diagnosed with early breast cancer similarily demonstrated that differences in the gut microbiome affected tumor aggressiveness, prognosis, and therapy response^6^, supporting the notion that the gut microbiome can systemically impact breast cancer outcomes.

The mechanism by which changes in the gut microbiome impact breast tumor metastasis remains unknown. Dysbiosis can trigger systemic inflammation which is associated with increased metastatic recurrence in breast cancer patients^7^. We previously found that dysbiosis elevates levels of CCL2 in mammary tissue prior to tumor initiation. When increased levels of CCL2 precede breast cancer, there is an increased risk of developing breast cancer^8^ or metastatic disease after diagnosis^9^. Considering the heterogenous outcomes for HR^+^ breast cancer and our previous findings, we hypothesized that commensal dysbiosis changes the immunological tone of the pre-malignant mammary tissue, and that the cellular and molecular changes that arise in response to dysbiosis enhance early metastatic spread when a tumor is present. Here, we sought to investigate how commensal dysbiosis-induced pro-inflammatory mediators in the mammary tissue influence dissemination of HR^+^ tumor cells.

Recently, McKee *et al*. demonstrated that antibiotic treatment enhances mast cell accumulation into mammary tumors, increasing tumor fibrosis and growth kinetics^10^. Mast cells are evolutionarily conserved innate immune cells that are highly attuned to the tissue environment of the organs they reside in. Mast cells mediate stromal rearrangement needed for the development of mammary ducts during mammary tissue development and homeostasis^11^, suggesting a role for the maintenance of mammary tissue homeostasis. However, mast cells are also capable of inducing endothelial cell tube formation^12^, fibrosis^13^ and inflammation^14^ in non-cancer disease models. In the context of cancer, these conditions associate with enhanced tumor dissemination. Despite the role of mast cells in mammary gland development and homeostasis, it is completely unknown whether mammary tissue-associated mast cells impact breast cancer metastasis.

In this study, we demonstrate that commensal dysbiosis enhances accumulation of mast cells in the mammary tissue of non tumor-bearing mice, in response to elevated mammary tissue CCL2 levels. In the presence of a tumor, pre-existing commensal dysbiosis results in tissue-infiltrating mast cells becoming pro-fibrogenic, culminating in increased tissue collagen levels and the promotion of early metastatic dissemination of HR^+^ tumors. We confirm that increased mast cell abundance correlates with high collagen levels in mammary tissues of women diagnosed with HR^+^ breast cancer. To our knowledge, this is the first study revealing that commensal microbiome-mediated accumulation and phenotypic changes in mammary tissue-associated mast cells influence the metastatic spread of HR^+^ tumor cells.

## Results

### Commensal dysbiosis drives accumulation of mast cells in mammary tissues

Previously, we demonstrated that the gut microbiome was a distal host-intrinsic mediator of breast tumor metastasis. When commensal dysbiosis was established prior to breast tumor initiation, metastatic dissemination of HR^+^ tumors to the axillary lymph node, blood, and lungs were significantly increased^5^. Importantly, dissemination of HR^+^ tumor cells did not occur due to differences in tumor growth, as tumor volumes were unchanged in response to dysbiosis^5^. This led us to hypothesize that microenvironmental changes arising in response to dysbiosis prime the mammary tissue to enhance metastatic dissemination. Commensal dysbiosis elevates mammary tissue levels of chemokines CCL2 and CXCL10 in non-tumor-bearing mice^5^. Although dysbiosis-induced tissue chemokines eventually return to baseline in non-tumor bearing mice, CXCL10 and CCL2 remain elevated in adjacent mammary tissues of HR^+^ tumor-bearing mice^5^. These findings suggest that CCL2 and/or CXCL10 could be tissue microenvironmental factors that contribute to dysbiosis-induced tumor dissemination of HR^+^ tumor cells.

To determine the corresponding cellular changes that arise in response to commensal dysbiosis and increased CCL2 and/or CXCL10 levels, we first analyzed the cellular composition of mammary tissues from non-tumor-bearing mice with or without commensal dysbiosis. Commensal dysbiosis was established with antibiotics prior to analysis of mammary tissues and mice were synchronized into estrus to normalize cellular changes occurring in response to the estrus cycle (Figure 1A), as previously described^5^. Macrophages and myeloid cells in the mammary gland increase risk of metastatic breast cancer^15, 16^, and can be recruited in response to CCL2^17^ and CXCL10^18^. Commensal dysbiosis alone did not increase myeloid populations in the mammary gland of non-tumor-bearing mice (Figure 1B), despite our previous finding that dysbiosis enhanced the accumulation of myeloid cells in adjacent mammary tissue and tumors during early and advanced stages of HR^+^ cancer^5^. Although CD4 T cells were slightly elevated in mammary tissues of non-tumor-bearing dysbiotic mice, the differences were not statistically significant and did not persist during tumor progression (Figure S1A). Similarly, no significant changes in the number of CD8 T cells, NK, NKT cells, and double negative T cells were observed at any point prior to or during tumor progression (Figure S1A-B).

**Figure 1:**
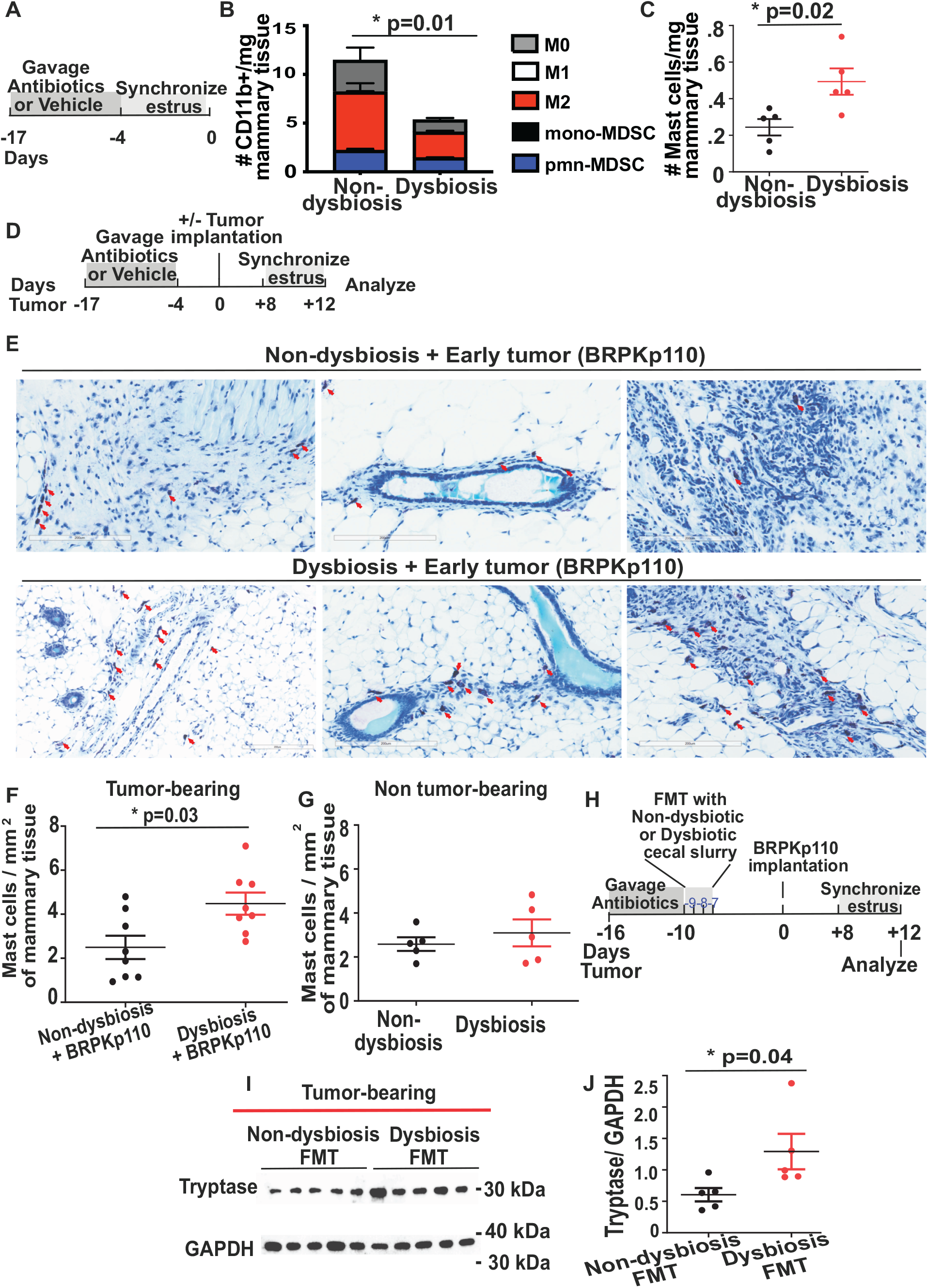
Mast cells accumulate in the mammary tissues of mice with commensal dysbiosis. **A,** Experimental scheme for B. C57BL/6 mice were orally gavaged daily with a broad-spectrum cocktail of antibiotics or an equal volume of water as a vehicle control for 14 days. Gavage was ceased for 4 days prior to sample collection. A modified Whitten effect was used to synchronize estrus prior to sample collection. **B,** C57BL/6 mice were treated as described in Fig. 1A. On day 0, myeloid cells were quantitated in disassociated mammary tissues. All populations were gated on live, singlet, CD45^+^CD11b^+^ cells. Numbers represent absolute numbers of cells quantitated using counting beads. M0 macrophages = F4/80^+^CD86^−^CD206^−^. M1 macrophages = F4/80^+^CD86^+^CD206^−^. M2 macrophages = F4/80^+^CD86^−^CD206^+^. Monocytic MDSC = Ly6C^hi^Ly6G^−^. Polymorphonuclear MDSC = Ly6C^mid^Ly6G^+^. **C,** Mast cells were quantitated in mammary tissues at day 0 using flow cytometry. **D,** Experimental scheme for E - G. Mammary tumor cells were orthotopically injected into the 4^th^ mammary fat pad of mice with or without commensal dysbiosis. Twelve days post tumor initiation, mammary tissues were evaluated. Non-tumor-bearing mice at the same point post antibiotic-induced dysbiosis were included for analysis. Mice were synchronized in estrus prior to sample collection. **E,** 12 days after BRPKp110 tumor implantation, mast cells in the adjacent mammary tissue were identified by toluidine blue staining. Mast cells are marked with red arrows. Representative of two independent experiments with 4-5 mice/group. Mast cells were enumerated by blindly counting within one mammary tissue whole mount section stained with toluidine blue from tumor-bearing mice **(F)** and non-tumor-bearing mice **(G)** followed by normalization to the entire mammary tissue area. **H,** Experimental scheme for the fecal microbiota transplantation and tumor implantation in I - J. C57BL/6 mice were orally gavaged for 7 days with a broad-spectrum cocktail of antibiotics. Immediately following the cessation of antibiotics, mice were orally gavaged for three consecutive days with cecal slurries collected from non-dysbiotic or dysbiotic mice collected at day 0, as depicted in Fig. 1A. Mice were rested for 7 days to allow for engraftment of transplanted microbial populations, followed by initiation of BRPKp110 tumors. Mice were evaluated 12 days post tumor initiation. **I and J,** protein levels of the mast cell marker tryptase were evaluated in mammary tissues by immunoblot. Protein levels of tryptase were quantitated using ImageJ and normalized to GAPDH **(J)**.

Mast cells are highly responsive to the microenvironment in which they reside, and in other non-cancer disease models have been shown to induce fibrosis^13^ and inflammation^14^, both of which are relevant to the changes that occur in the mammary tissue of tumor-bearing mice with pre-existing commensal dysbiosis^5^. Interestingly, mast cell numbers significantly increased in mammary tissue in response to commensal dysbiosis (Figure 1C and S1C). Importantly, changes in mast cell numbers mirrored the changes in mammary tissue levels of CCL2 and CXCL10. Specifically, mast cell numbers remained elevated in the adjacent mammary tissue of dysbiotic mice after tumor implantation (Figure 1 D-F), whereas a return to baseline levels was observed in the mammary tissues of non-tumor-bearing mice (Figure 1D and 1G). Tryptase, a serine protease predominantly produced by mast cells, serves as a specific mast cell marker in both animal and human studies^19^. In line with increased mast cell numbers observed in adjacent mammary tissues in tumor-bearing mice, significantly enhanced protein levels of tryptase were detected in mammary tissue lysates of adjacent mammary tissues from dysbiotic tumor-bearing mice when compared to levels in non-dysbiotic mice (Figures S1D-E). Mice bearing the highly metastatic PyMT tumor model^20^ also had increased mast cell abundance in the adjacent mammary tissue in response to commensal dysbiosis (Figure S1F-G), suggesting that the observed changes in mast cell numbers are not tumor model-specific.

The lung is a major reservoir of mast cells^21^, and a major site for breast tumor metastasis^1^. To evaluate the the effects of dysbiosis on mast cell abundance in lungs, flow cytometry was used to evaluate lungs from mice prior to and during tumor progression. Interestingly, the abundance of mast cells was increased both in response to dysbiosis in non-tumor-bearing mice (Figure S1H) and in tumor-bearing mice (Figure S1I). These results indicate that commensal dysbiosis increases the abundance of mast cells in lungs of mice with mammary tumors.

We have previously shown that fecal microbiota transplantation (FMT) of a dysbiotic microbiome enhances metastatic dissemination of HR^+^ breast tumors^5^. As an additional readout of dysbiosis-induced mast cell expansion, we next tested whether dysbiotic FMT also enhances numbers of mast cells in the mammary tissues of tumor-bearing mice. To do this, cecal contents from dysbiotic and non-dysbiotic donor mice were orally gavaged into antibiotic-sterilized recipient mice for 3 consecutive days (Figure 1H). After allowing a week for the transplanted microbiome to engraft in the recipient mice, tumor cells were implanted and mice were analyzed 12 days after (Figure 1H). Mice transplanted with a dysbiotic microflora had significantly increased tryptase protein levels in adjacent mammary tissues (Figure 1I-J). These results indicated that commensal dysbiosis was sufficient to enhance tryptase, and thus mast cells, in mammary tissues during HR^+^ tumor progression.

### Mast cells promote early dissemination of HR^+^ mammary tumor cells

We next asked whether mast cell activation influences the dissemination of HR^+^ breast tumor cells. To do this, mice were orally gavaged with the mast cell stabilizer ketotifen fumarate (10 mg/kg) once daily from days 1 to 5 following implantation of the poorly-metastatic HR^+^ tumor line BRPKp110 into the mammary fat pad (Figure 2A). Ketotifen treatment significantly reduced dysbiosis-mediated tumor dissemination into the lungs (Figure 2B and S2A) at advanced stages of tumor progression. Importantly, ketotifen treatment did not affect the primary BRPKp110 tumor growth (Figure S2B). Ketotifen treatment also significantly reduced the lung dissemination of the highly-metastatic HR^+^ breast tumor cell line PyMT (Figure 2C and S2C). Because the treatment of ketotifen treatment reduced the PyMT primary tumor volumes (Figure S2D), the numbers of PyMT tumor cells in the lungs were normalized to final tumor burden (Figure 2C) to account for variation in tumor growth. Interestingly, ketotifen treatment reduced tumor dissemination in non-dysbiotic mice bearing PyMT tumors (Figure 2C), suggesting that mast cell-mediated tumor dissemination may also be influenced by tumor-intrinsic factors. Altogether, these data suggest that mast cell activation contributes to dysbiosis-induced metastatic dissemination of HR^+^ breast tumors.

**Figure 2:**
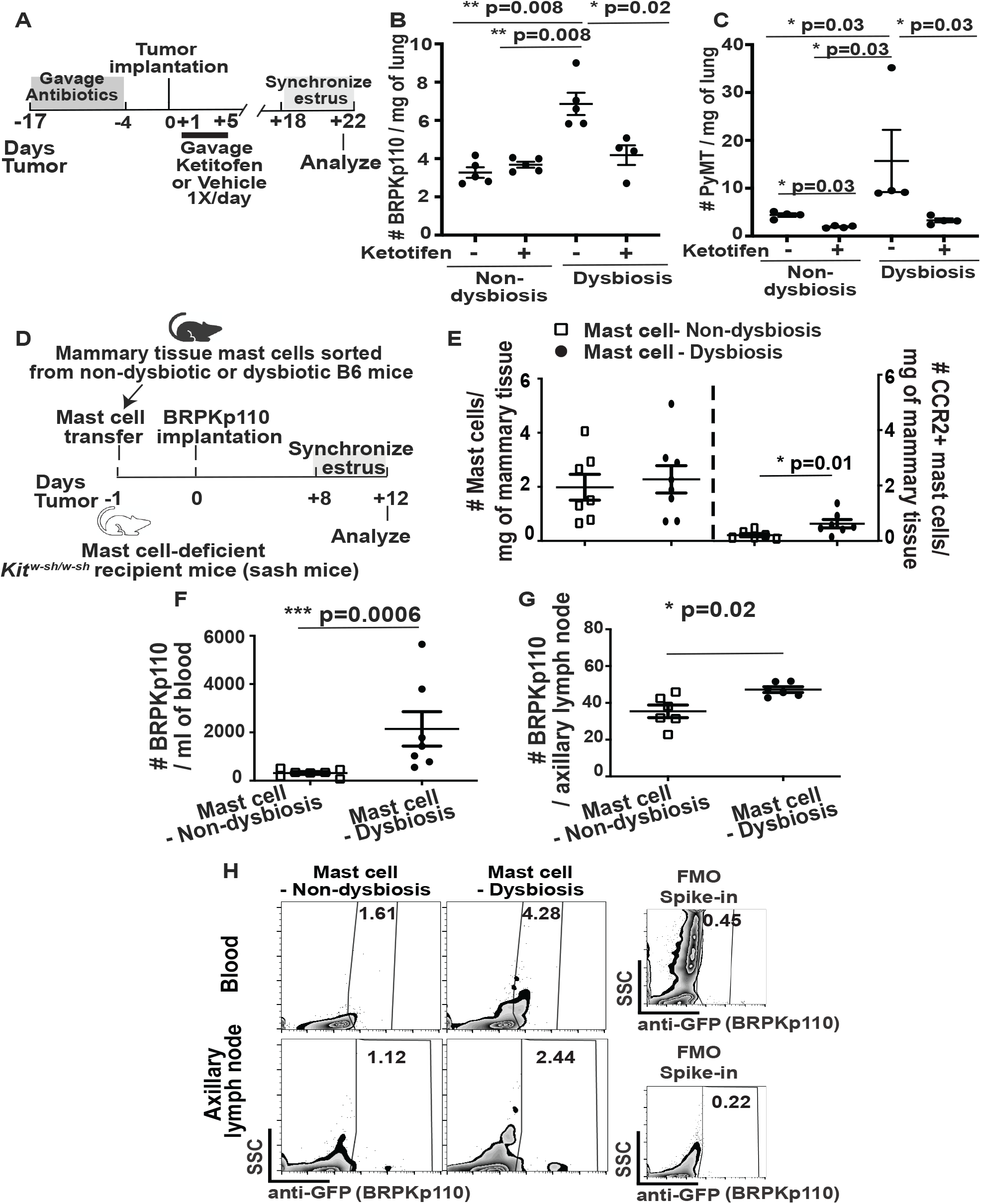
Mast cells promote dissemination of HR^+^ tumor cells in response to commensal dysbiosis. **A,** Experimental scheme for ketotifen treatment in B and C. Tumor-bearing dysbiotic or non-dysbiotic mice were gavaged with ketotifen or vehicle daily starting 1-day post tumor initiation and continuing to day 5 post-tumor. Mice were evaluated 22 days post tumor initiation. **B and C,** Lungs from mice bearing advanced BRPKp110 or PyMT tumors were evaluated for disseminated GFP^+^ BRPKp110 **(B)** or luciferase^+^ PyMT **(C)** tumor cells 22 days post tumor initiation using flow cytometry. Numbers of disseminated PyMT were normalized to the final tumor volumes. **D,** Experimental scheme for E - G. Equal numbers of mammary tissue mast cells sorted from non-dysbiotic or dysbiotic C57BL/6 mice from day 0 were transferred into the inguinal mammary fat pad of mast cell-deficient *Kit^w-sh/w-sh^* mice. After 18 hours post transfer, BRPKp110 tumor cells were injected into the same mammary fat pad of *Kit^w-sh/w-sh^* recipient mice. Twelve days post tumor implantation, mice were evaluated for disseminated tumor cells and transferred mast cells within the mammary tissues. **E,** Flow cytometry was utilized to quantitate absolute numbers of mast cells and CCR2^+^ mast cells remaining post transfer in the adjacent mammary tissues of tumor-bearing *Kit^w-sh/w-sh^* mice. Numbers represent mast cells/gram of mammary tissue. **F - G,** Dissemination of GFP^+^ BRPKp110 tumor cells was quantitated from the peripheral blood **(F)**, and tumor-draining axillary lymph nodes **(G)** by flow cytometry 12 days post tumor initiation in *Kit^w-sh/w-sh^* mice. **H,** Representative density flow cytometry plots showing GFP^+^ BRPKp110 tumor cells in the peripheral blood and tumor-draining axillary lymph nodes of *Kit^w-sh/w-sh^* mice. FMO spike-in represents a gating control of blood or lymph nodes spiked with BRPKp110 tumor cells, minus the anti-GFP antibody. Numbers represent the percentage of gated GFP^+^ tumor cells of CD45 negative cells. Cumulative of two independent experiments with 3 mice/group. Each symbol represents an individual mouse, and statistical significance was determined by two-tailed Mann-Whitney *U* test.

Next, we examined if mast cells in mammary tissue of dysbiotic animals were sufficient to induce metastatic dissemination. Mast cells were sorted from mammary tissues of non-tumor-bearing dysbiotic and non-dysbiotic mice and equal numbers were orthotopically transferred into the inguinal mammary fat pad of mast cell-deficient *Kit^w-sh/w-sh^* mice (Sash mice)^22^. The following day, BRPKp110 tumor cells were injected into the same mammary fat pad (Figure 2D). Using flow cytometry, we confirmed that equivalent numbers of mast cells remained in the mammary tissues of dysbiotic and non-dysbiotic mice 12 days following tumor initiation (Figure 2E), suggesting that both groups of sorted mast cells were equally capable of engrafting into the mammary tissue of Sash mice. Sash mice that received mast cells sorted from mammary tissues from dysbiotic mice had significantly increased dissemination of tumor cells in the blood and the axillary lymph nodes (Figure 2F-H), when compared to the Sash mice that received mast cells from the mammary tissues of non-dysbiotic mice (Figure 2F-H). To test whether dysbiosis can induce tumor dissemination in the absence of mast cells, we subjected Sash mice to the transfer of a dysbiotic microbiome using FMT prior to tumor initiation (Figure S2E). Dysbiotic Sash mice showed comparable tumor growth (Figure S2F) and reduced dissemination of tumor cells (Figure S2G-H) compared to that observed in wild-type mice transplanted with a dysbiotic microbiome (Figure S2G-H). Collectively, these data indicate that mast cells are required for dysbiosis-induced metastatic dissemination of HR^+^ tumors. Furthermore, these results suggest that mammary tissue mast cells are functionally changed in response to commensal dysbiosis, and that these functional changes increase dissemination of HR^+^ tumor cells.

### CCL2-CCR2 signaling drives dysbiosis-induced mast cell accumulation in mammary tissue and subsequent early dissemination of HR^+^ tumor cells

Changes in mast cell numbers mirrored changes in chemokine levels of CCL2 and CXCL10^5^, leading us to hypothesize that mast cells are accumulating in response to the elevation in CCL2 and/or CXCL10. To test this hypothesis, we first measured CCR2 and CXCR3 levels on mast cells in the mammary tissue of dysbiotic and non-dysbiotic mice, with the assumption that expression of CCR2 or CXCR3 corresponds with accumulation in response to each respective ligand, CCL2 or CXCL10. Using flow cytometry, we found that dysbiotic mice had increased numbers of CCR2^+^ mast cells in mammary tissue when compared with non-dysbiotic mice whereas no differences were observed for CXCR3-expressing mast cells (Figure 3A). Notably, we also observed enrichment of a CCR2^+^ mast cell population in Sash mice that received mast cells from dysbiotic mice (Figure 2E). These data suggest that mast cells are accumulating in response to CCL2, and that CCR2 expression is a stable signature of mammary tissue mast cells from dysbiotic mice.

**Figure 3:**
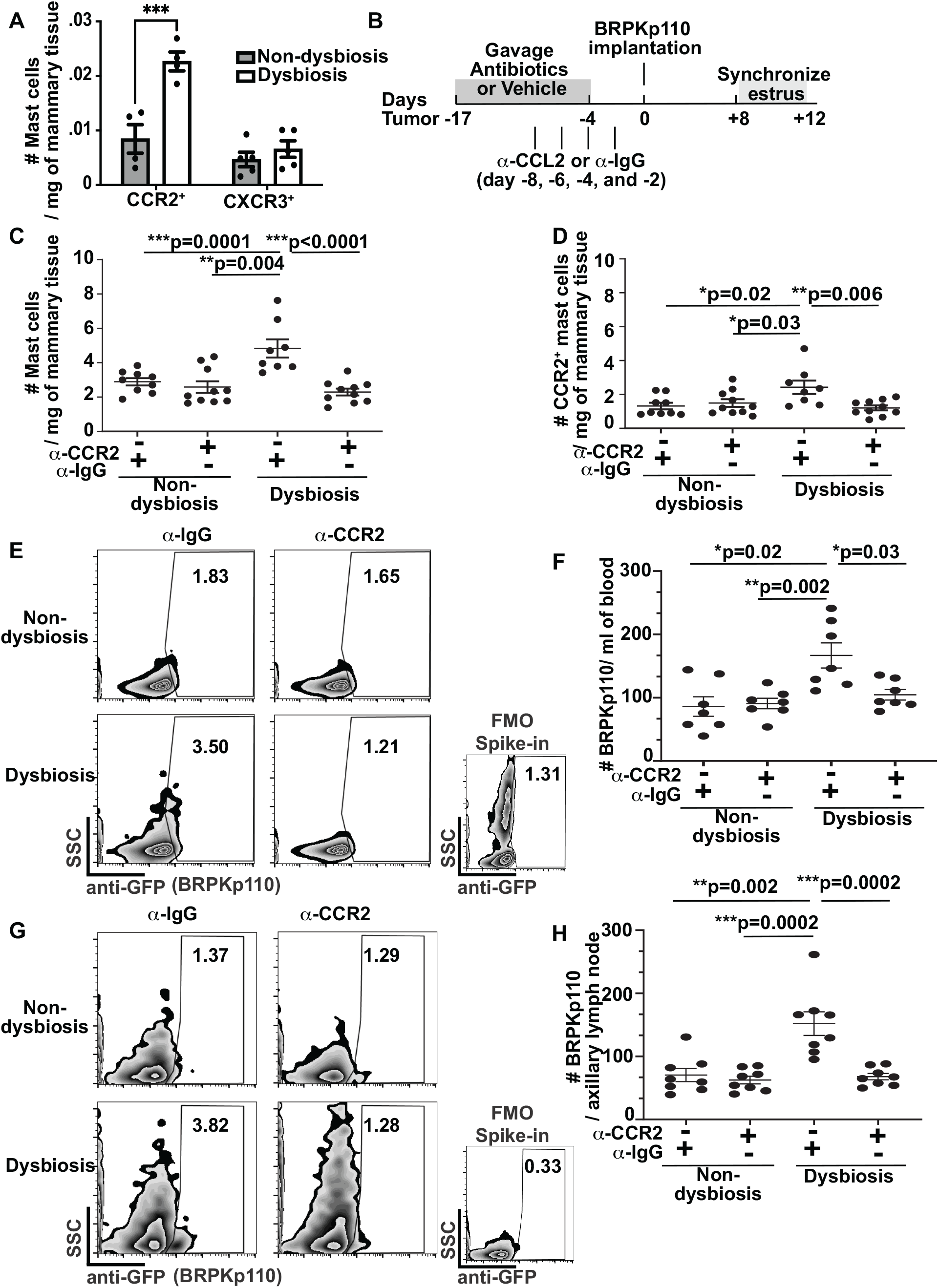
CCL2-CCR2 signaling leads to dysbiosis-induced mast cell accumulation into mammary tissue and subsequent early dissemination of HR^+^ tumor cells. **A,** Mammary tissue mast cells were evaluated in dysbiotic and non-dysbiotic tumor-bearing mice 12 days post tumor initiation. CCR2^+^ and CXCR3^+^ mammary tissue mast cells were quantitated using flow cytometry. **B,** Experimental design for CCL2 blockade prior to tumor initiation (C to H). Mice were intraperitoneally injected with 4 doses of anti-CCL2 or an isotype-matched control IgG prior to tumor initiation on day −8, day −6, day −4 and day −2. Tumors were initiated on day 0. **C to H**, 12 days post tumor initiation, mammary tissues **(C and D)** were evaluated and tumor dissemination **(E to H)** was quantified. Mast cells **(C)** and CCR2^+^ mast cells **(D)** from adjacent mammary tissues were quantitated using flow cytometry. GFP^+^ BRPKp110 tumor cells were quantitated from the peripheral blood (representative flow plots in **E**, and tumor quantification in **F**) and tumor-draining axillary lymph nodes (representative flow plots in **G**, and tumor quantification in **H**). For flow plots, numbers represent the frequency of GFP^+^ tumor cells from CD45 negative cells. Each symbol represents an individual mouse, and statistical significance was determined by two-tailed Mann-Whitney *U* test.

To test whether mast cells are accumulating in the mammary tissue in response to CCL2, we neutralized CCL2 using an antibody based approach on 8, 6, 4, and 2 days prior to tumor initiation (Figure 3B). Neutralization of CCL2 significantly reduced the abundance of total (Figure 3C and S3A) and CCR2^+^ (Figure 3D and S3B) mast cells infiltrating into adjacent mammary tissues of dysbiotic mice. Reduced mast cell accumulation was not due to changes in tumor growth, as overall tumor burden was similar across groups (Figure S3C). Corresponding with the reduction of mast cells during CCL2 blockade, tumor dissemination into the peripheral blood (Figure 3E-F) and tumor-draining lymph nodes (Figure 3G-H) was significantly reduced only in dysbiotic mice. Together, these results indicate that CCL2 is required for commensal dysbiosis-induced accumulation of mammary tissue mast cells and early dissemination of HR^+^ tumor cells.

### Mast cells are involved in commensal dysbiosis-mediated fibroblast activation in adjacent mammary tissues

Mast cells contribute to fibroblast activation and tissue fibrosis in various organs in both humans and animals^13^. Activated fibroblasts, also called myofibroblasts, are involved in extracellular matrix (ECM) synthesis and remodeling, creating a conduit for tumor cell egress out of the primary tissue and dissemination into peripheral organs^23^. To determine whether mast cells affected fibroblast activation and matrix remodeling, we first analyzed mammary tissue fibroblasts from mice prior to and during early tumor progression (Figure 4A). Collagen I^+^ and smooth muscle actin (SMA)^+^ were evaluated as markers of activation on fibroblasts using flow cytometry (Figure S4A). Commensal dysbiosis enhanced fibroblast expansion, resulting in increased numbers of activated fibroblasts in the normal non-tumor-bearing mammary tissue (Figure 4B). Similar to the kinetics of mast cell infiltration into the mammary tissue, the numbers of activated fibroblasts diminished in the absence of a tumor (Figure 4C). Importantly, in the presence of a tumor, there was a significant expansion of activated fibroblasts in adjacent tissue in response to commensal dysbiosis (Figure 4D). As expected, collagen I protein levels in the adjacent mammary tissues of tumor-bearing dysbiotic mice bearing BRPKp110 (Figures 4E, S4B) or PyMT (Figures 4F, S4C) tumors were similarly increased, when compared with non-dysbiotic mice with equivalent tumor burden.

**Figure 4:**
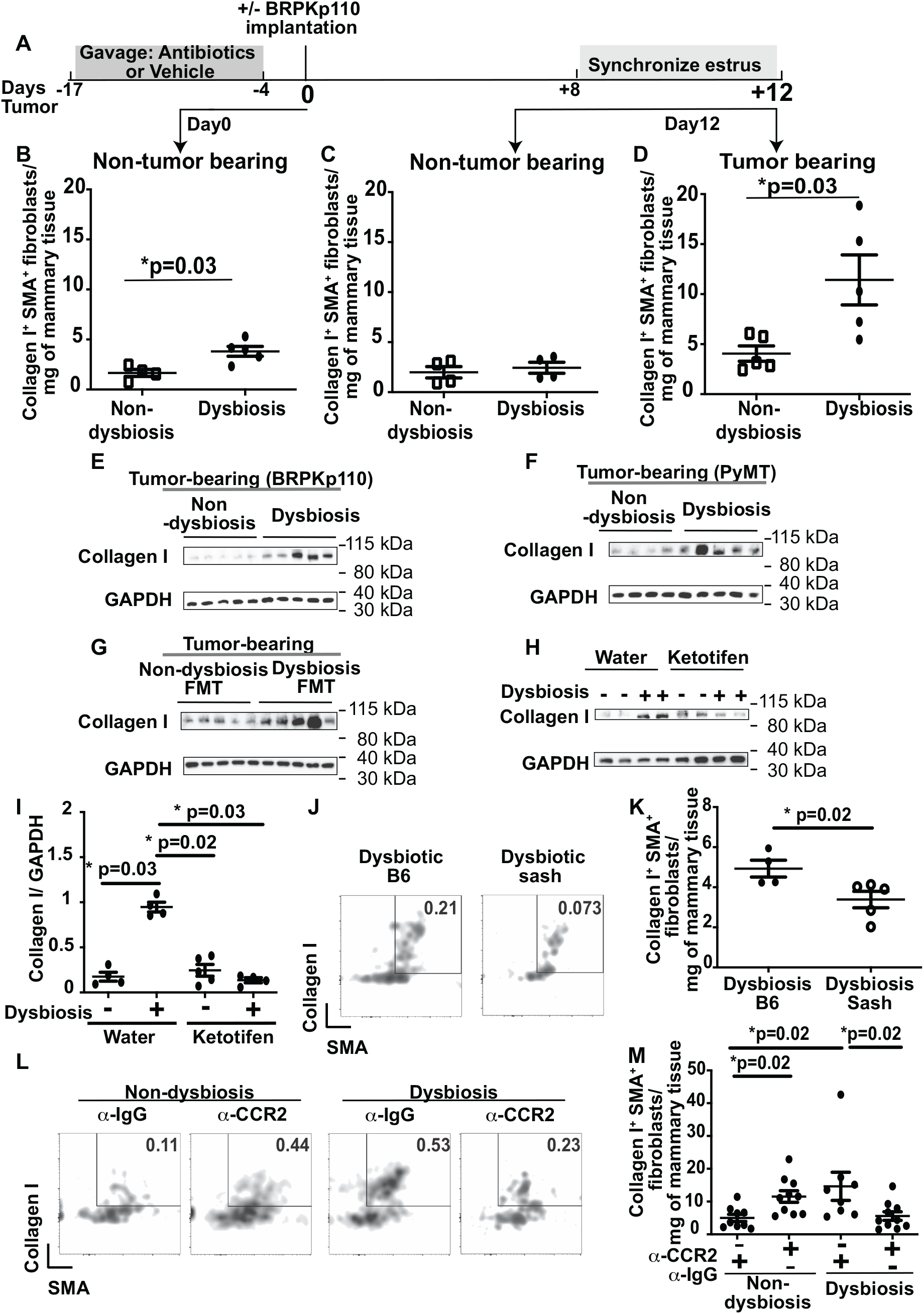
Commensal dysbiosis-induced mast cells activate tissue fibroblasts to produce collagen I. **A,** Experimental scheme for B - D. Samples were collected at day 0 for B and +12 for C and D. **B,** Numbers of collagen I^+^ SMA^+^ fibroblasts in the mammary tissues of non-tumor-bearing dysbiotic or non-dysbiotic mice at day 0. **C and D,** Quantification of collagen I^+^ SMA^+^ fibroblasts in the mammary tissues from non-tumor-bearing **(C)** or tumor-bearing **(D)** mice on day + 12. Numbers are represented per gram of mammary tissue. **E and F,** Protein levels of collagen I in the adjacent mammary tissues of mice bearing BRPKp110 **(E)** or PyMT **(F)** mammary tumors after 12 days. Collagen I was measured using immunoblot. GAPDH was included as a protein loading control. **G,** Mice received a fecal transplant of non-dysbiotic or dysbiotic cecal slurries prior to tumor initiation, as in Figure 1G. Levels of collagen I protein in the mammary tissue were evaluated 12 days post tumor implantation. **H and I,** Tumor-bearing non-dysbiotic or dysbiotic mice were treated with the mast cell stabilizer ketotifen or water as a control. Collagen I levels in the mammary tissue were evaluated by immunoblot 12 days post tumor implantation **(H)** and quantified based on GAPDH levels using ImageJ **(I)**. **J and K,** Dysbiosis was established in wild-type C57BL/6 mice (B6) or mast cell-deficient *Kit^w-sh/w-sh^* mice (Sash) as depicted in Fig. S1J. Collagen I^+^ SMA^+^ fibroblasts in the mammary tissues were quantified using flow cytometry 12 days post tumor implantation. **(J)** Representative density plots depicting the percentage of collagen I^+^ SMA^+^ fibroblasts of total live cells. **(K)** Quantitation of total fibroblasts per gram of mammary tissue using flow cytometry. **L and M,** anti-CCL2 antibody or control IgG were intraperitoneally injected into non-dysbiotic and dysbiotic mice prior to tumor implantation as depicted in Fig. 2B. At day +12, collagen I^+^ SMA^+^ fibroblasts in the mammary tissues were quantitated using flow cytometry. **(L)** Representative density plots depicting the percentage of collagen I^+^ SMA^+^ fibroblasts of total live cells. **(M)** Quantitation of total fibroblasts per gram of mammary tissue using flow cytometry. For all flow cytometric analysis of fibroblasts, fibroblasts were identified as CD45^+^ CD31^−^ gp38^+^ PDGFR-α^+^ cells. Each symbol represents an individual mouse, and statistical significance was determined by two-tailed Mann-Whitney *U* test.

Using the FMT approach, we found that transfer of a dysbiotic microflora was sufficient to induce collagen I production in the adjacent mammary tissues of BRPKp110 tumor-bearing mice (Figures 4G, S4D). Interestingly, similar to changes in adjacent mammary tissues, tumors from dysbiotic mice also had increased collagen I protein levels (Figure S4E), with a significant linear correlation between the protein levels of collagen I in the mammary tissue and the protein levels of collagen I in the tumor (Figure S4F). However, this association was not observed in tumors from non-dysbiotic mice (Figure S4G). Fibroblasts in mammary tumors are recruited from the adjacent mammary tissue environment^24^. In line with these findings, our data indicate that commensal dysbiosis leads to increased collagen levels in mammary tissue and tumors.

We next evaluated whether mast cell accumulation into the tissue environment affects activation of tissue fibroblasts and deposition of collagen. Dysbiosis was initiated and mice were treated with the mast cell stabilizer ketotifen, as depicted in Figure 2A. Ketotifen significantly reduced the accumulation and activation of fibroblasts in the adjacent mammary tissue of tumor-bearing mice with commensal dysbiosis (Figure S4H). Protein levels of collagen I in the adjacent mammary tissue of dysbiotic mice treated with ketotifen were correspondingly reduced (Figure 4H-I). Next, we compared whether fibroblasts would be activated in response to commensal dysbiosis and HR^+^ tumors in the absence of mast cells. To do this, dysbiosis was initiated in wild-type and mast cell-deficient *Kit^w-sh/w-sh^* Sash mice, similar to Figure S2E. Compared to C57BL/6 mice, the Sash mice had reduced numbers of collagen I^+^ SMA^+^ fibroblasts in the adjacent mammary tissues (Figure 4J-K). Reducing mammary tissue mast cell accumulation by neutralizing CCL2 also reduced the abundance and activation of fibroblasts in the adjacent mammary tissue of dysbiotic mice (Figure 4L-M). Together, these results suggest that commensal dysbiosis-induced mammary tissue mast cells initiate fibroblast activation and shift the tissue millieu towards a pro-fibrogenic microenvironment.

### Increased PDGF-B^+^ mast cell numbers in dysbiotic mice correspond with increased PDGFR-a expression on fibroblasts

Transfer of mast cells into Sash mice (Figures 2E-H and Figure S2E-H) suggested that mast cells from dysbiotic mice were also phenotypically distinct from those found in mammary tissues of non-dysbiotic mice. To address this hypothesis, we first evaluated expression levels of genes related to pro-fibrogenic pathways from mast cells sorted from mammary tissues of dysbiotic and non-dysbiotic mice. Using real-time PCR, we found that the gene expression of *Pdgfb* was increased in mammary tissue mast cells from tumor-bearing dysbiotic mice, while other pro-fibrogenic mediators, including *Mcpt4, Mcpt5, Mcpt2, Pdgfa*, and *Tgfb1*, remained unchanged (Figure S5A). The mast cell marker *Cpa3* remained comparable between the two groups (Figure S5A). Interestingly, tumor-associated mast cells from dysbiotic mice showed increased *Mcpt2*, *Pdgfb*, and *Cpa3* compared to that observed in tumor-associated mast cells from non-dysbiotic mice (Figure S5B), supporting the idea that the phenotype of mast cells is greatly influenced by the local microenvironment.

Pro-fibrogenic mediator platelet-derived growth factor B (PDGF-B) is known to be involved in mast cell-mediated activation of fibroblasts^25^. Indeed, numbers of PDGF-B-expressing mast cells in mammary tissues of dysbiotic mice were significantly increased compared with numbers of PDGF-B^+^ mast cells in mammary tissues of non-dysbiotic mice (Figure 5A). Importantly, mammary tissues of dysbiotic mice retained significant numbers of PDGF-B^+^ mast cells during early tumor progression (Figure 5B), and these changes corresponded with a significant increase in the levels of the receptor for PDGF-B, platelet-derived growth factor receptor a (PDGFR-α), on mammary tissue fibroblasts from dysbiotic mice prior to (Figure 5C) and during tumor progression (Figure 5D). These data suggest that mammary tissue mast cells, triggered in response to commensal dysbiosis, may promote fibrosis through the PDGF signaling pathway.

**Figure 5:**
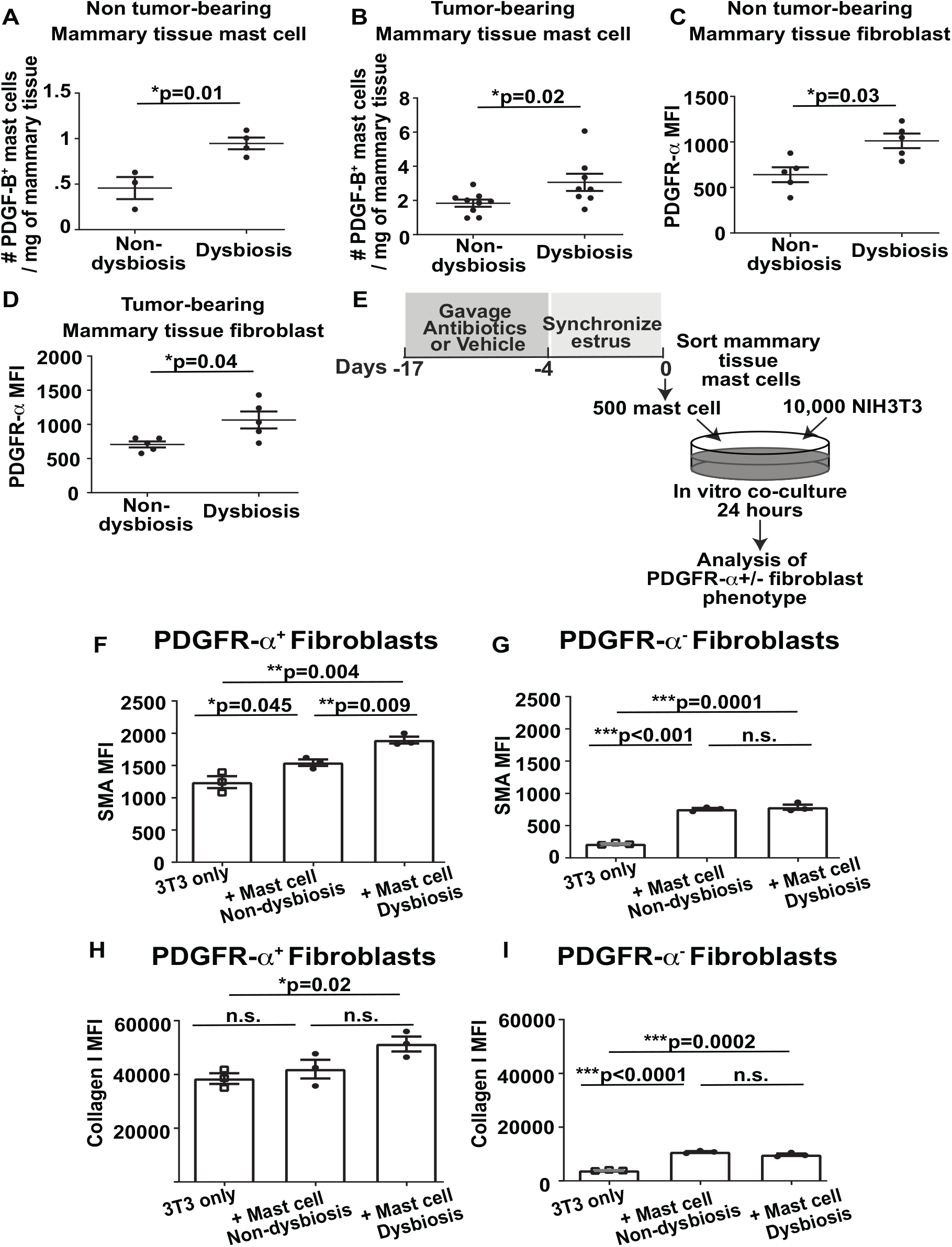
During early tumor progression, commensal dysbiosis increases expression of PDGF-B^+^ on mast cells with corresponding increases in PDGF receptor expression on fibroblasts. **A and B,** Numbers of PDGF-B^+^ mast cells in the mammary tissues from non-tumor-bearing **(A)** and tumor-bearing mice **(B).** Mast cells were quantitated using flow cytometry on mammary tissues disassociated into single cell suspensions on day +12, as depicted in Figure 4A. **C and D**, Evaluation of PDGFR-α staining intensity (mean fluorescent intensity - MFI) in mammary tissue fibroblasts from non-tumor-bearing **(C)** and tumor-bearing mice **(D)**. Each symbol represents an individual mouse, and statistical significance was determined by two-tailed Mann-Whitney *U* test. **E to I,** NIH-3T3 fibroblasts were co-cultured with mast cells that were sorted from mammary tissues of non-dysbiotic or dysbiotic mice at day 0, as depicted in **E**. Sorted mast cells were plated with NIH-3T3 fibroblasts at a ratio of 1:20. In some cultures, NIH-3T3 fibroblasts were plated without mast cells as negative controls. After 24 hours co-culture, the intensity of SMA and collagen I were measured using flow cytometry of either CD45^−^ PDGFR-α^+^ **(F and G)** or CD45^−^ PDGFR-α^−^ **(H and I)** fibroblast populations in each well, using mean fluorescent intensity (MFI) as the readout. Each symbol represents an experimental replicate, and statistical significance was determined by two-tailed unpaired t-test.

To directly test whether mammary tissue mast cells from dysbiotic mice can promote fibrosis, mast cells sorted from mammary tissues of dysbiotic or non-dysbiotic mice were cocultured with NIH-3T3 fibroblasts for 24 hours (Figure 5E). The following day, activation of PDGFR-α-positive and PDGFR-a-negative fibroblasts was evaluated by measuring expression levels of SMA and collagen I using flow cytometry. Mast cells sorted from mammary tissues of dysbiotic mice induced significantly greater levels of SMA and increased levels of collagen I in the PDGFR-α^+^ population of fibroblasts (Figure 5F and 5H). Although the collagen levels between PDGFR-α^+^ fibroblasts cultured with dysbiotic versus non-dysbiotic mast cells were not significantly different, collagen I levels were significantly increased when comparing NIH-3T3 fibroblasts cultured alone versus those cultured with mast cells from dysbiotic mice. On the other hand, mast cells from dysbiotic mice induced similar activation of PDGFR-α^−^ fibroblasts (Figure 5G and 5I), suggesting that PDGF signaling may be involved in mast cell-mediated fibroblast activation.

To determine how global protein levels of PDGF-B in the mammary tissue are affected by dysbiosis and HR^+^ tumors, PDGF-B protein levels were measured using immunoblot. Dysbiotic tumor-bearing mice had enhanced PDGF-B levels compared with mammary tissues from non-dysbiotic mice (Figures S6A-B). Additionally, a dysbiotic microflora was sufficient to induce PDGF-B protein levels (Figure S6C-E). Importantly, tumor-bearing Sash mice had significantly lower levels of PDGF-B in response to FMT-induced dysbiosis (Figure S6F-G). As expected, mast cell deficiency also reduced levels of tryptase in the mammary tissue (Figure S6F and S6H).

Transforming growth factor beta-1 (TGF-β1) is another potent pro-fibrogenic and prometastatic mediator in multiple disease models^26^. However, total protein levels of TGF-β1 in the adjacent mammary tissues and tumors of dysbiotic and non-dysbiotic mice were comparable (Figure S6I-J). These results suggest that TGF-β1 is not a major mediator of dysbiosis-mediated fibroblast activation and subsequent HR^+^ tumor dissemination. Altogether, these results implicate mast cell-derived PDGF signaling as a major mediator of fibroblast activation and establishment of a pro-metastatic microenvironment.

### The abundance of mammary tissue-associated mast cells is positively correlated with fibrosis in HR^+^ breast cancer patients

To determine the clinical relevance of mast cell-mediated fibroblast activation with breast cancer, we evaluated tissue sections from a small archival cohort of HR^+^ breast cancer patients. Consecutive tissue sections were cut from formalin fixed and paraffin embedded tissue specimens containing tumor-adjacent mammary tissue. Tissue sections were stained with PicroSirius Red to measure collagen levels and toluidine blue to enumerate mast cells. In line with our murine studies, collagen levels were positively and significantly correlated with mast cell numbers (Figure 6A-B), suggesting that mast cell abundance corresponds with collagen levels in HR^+^ breast cancer patients.

**Figure 6:**
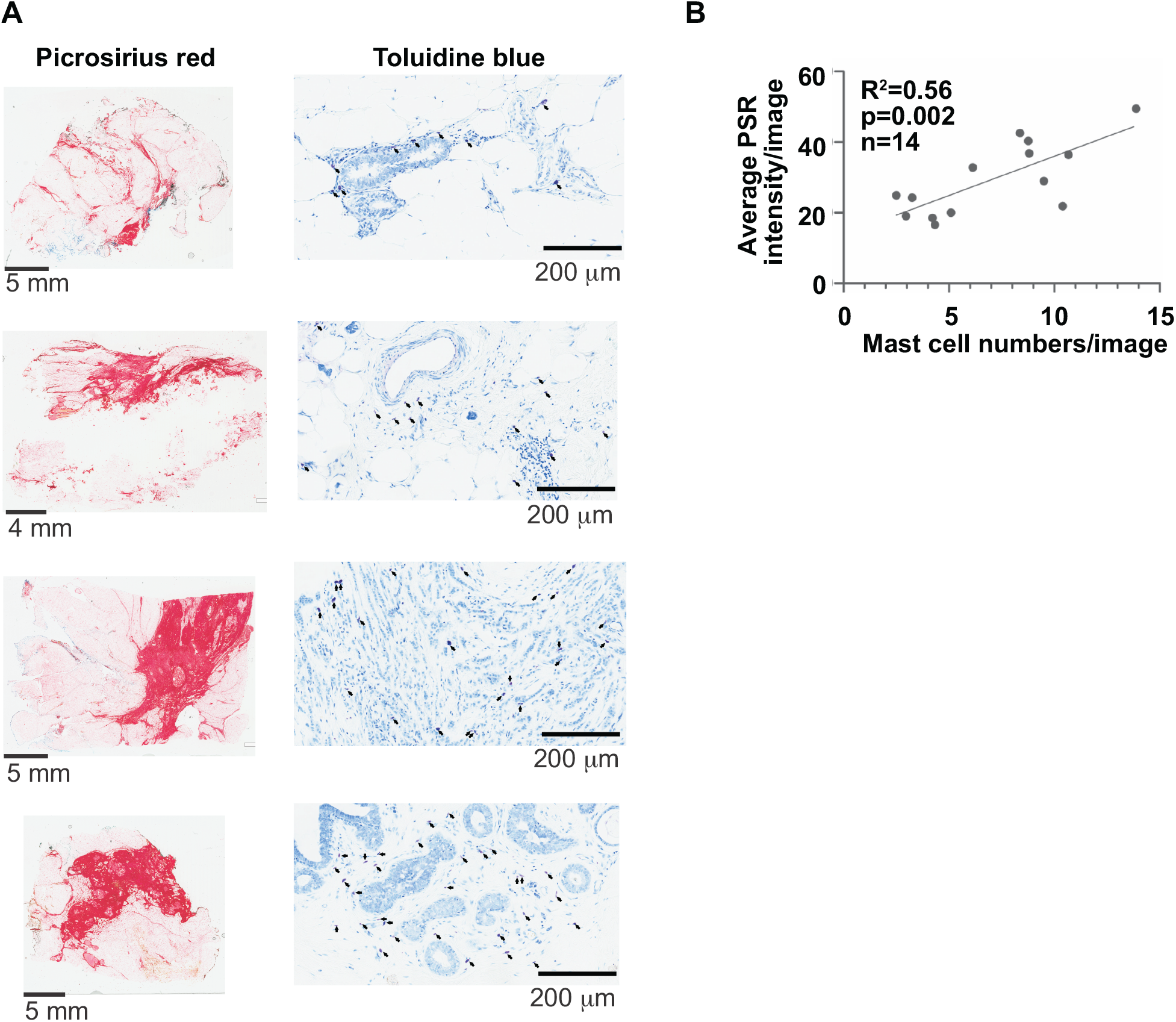
Mast cell abundance is positively correlated with collagen levels in breast tissues from HR^+^ breast cancer patients. **A,** Formalin-fixed paraffin-embedded (FFPE) breast tissues from 14 individuals previously diagnosed with HR^+^ breast cancer were sectioned and stained with PicroSirus Red and toluidine blue to evaluate collagen levels and to enumerate mast cells, respectively. Representative images are shown. 1X and 20X for PicroSirus Red (scale bar= 4 mm or 5 mm as indicated in the figure) and toluidine blue (scale bar= 200 μm), respectively. **B,** Linear correlation of PicroSirus Red staining intensity versus mast cell number/image. PicroSirus Red intensity was calculated using ImageJ. Mast cells were quantitated by averaging the total mast cells counted for 15 - 20 randomly acquired 10X microscopic images, encompassing the entire tissue section. Consecutive tissue sections were evaluated for each respective stain in a blinded manner. Each symbol represents an individual human sample, and the correlation was determined by linear regression.

### Mast cell signatures and myofibroblastic genes are enriched in the mammary tissues of women susceptible to breast cancer

Finally, we wanted to investigate whether the presence of mast cells and fibroblast activation may be detected in the mammary tissues of women prior to tumor diagnosis. To address this, we analyzed the transcriptomic sequencing data available from the Susan G. Komen Tissue Bank at Indiana University Simon Comprehensive Cancer Center. Normal breast tissues from either women prior to their breast cancer diagnosis (susceptible normal^27^) or age-matched healthy women were obtained. Patient characteristics are detailed in Table S2. Using principal component analysis (PCA), we found clustering of genes expressed in susceptible normal tissue while healthy donors showed a more heterogenous gene expression pattern (Figure 7A). Using gene set enrichment analysis, we found that mast cell gene signatures^28^ and PDGF-B (Table S1) were increased in the susceptible donors, while the breast cancer-associated myofibroblast genes^29^ (Table S1) were significantly enriched (Figure 7B-C). Collectively, these results indicate that mast cell-mediated fibroblast activation may arise before diagnosis with breast cancer.

**Figure 7:**
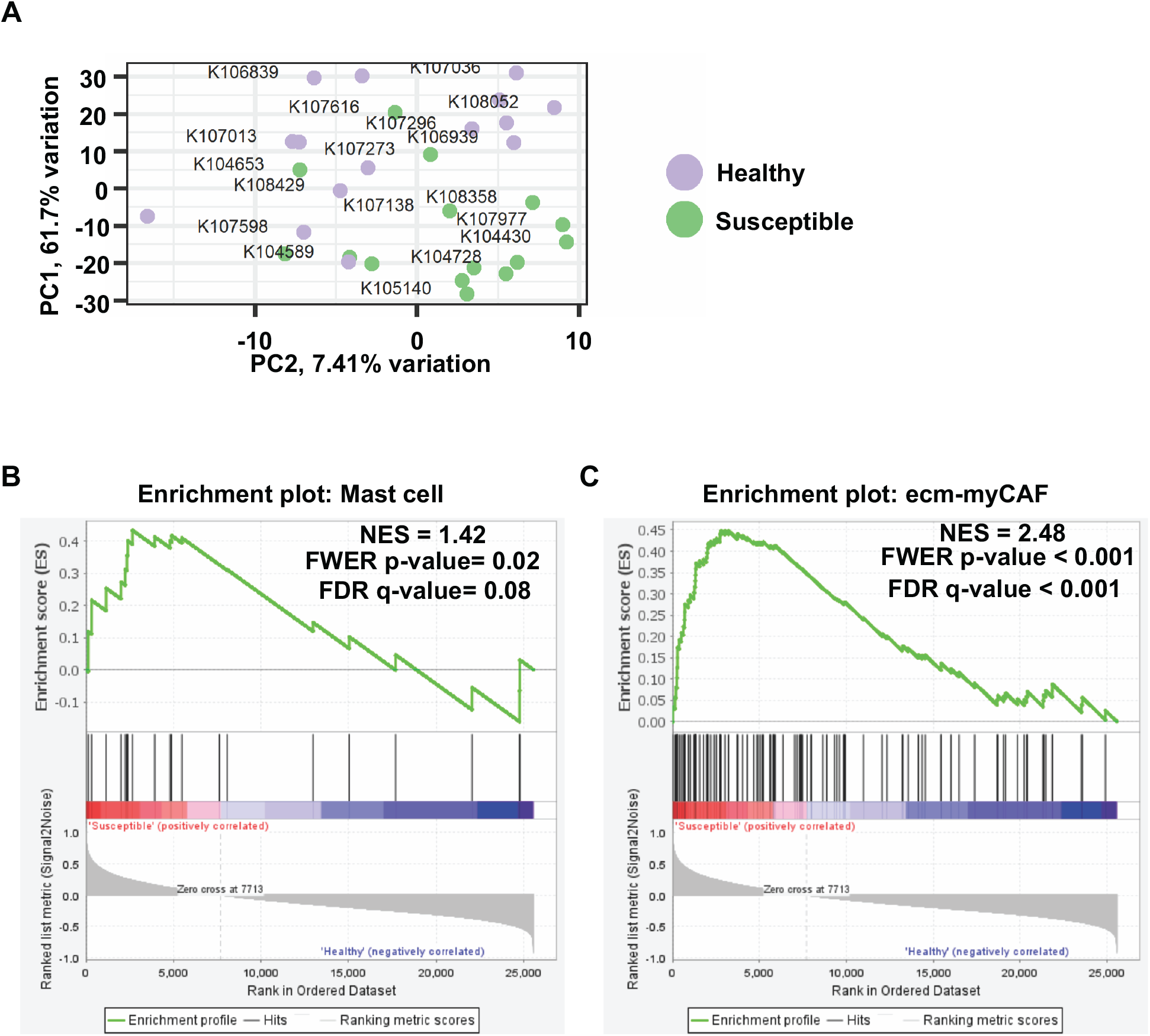
Normal breast tissues from women who later developed breast cancer have transcriptomic enrichment of mast cell and myofibroblastic gene signatures. **A,** Principal component analysis (PCA) of the age-matched healthy donors and women who later went on to develop breast cancer (susceptible). **B and C,** Gene set enrichment analysis (GSEA) for mast cell **(B)** and myofibroblastic **(C)** genes was performed on the RNA-seq data. N=15 donated normal breast tissue samples per group. NES: Normalized enrichment score; FDR: False discovery rate; FWER: Familywise-error rate.

## Discussion

Here, we demonstrate that mammary tissue mast cells enhance metastatic dissemination of HR^+^ tumors in response to gut commensal dysbiosis. The role of mast cells in breast cancer has been controversial. Previous studies have primarily focused on tumor- and lymph-node associated mast cells. Some studies identified a positive association with outcome^30, 31^ whereas others associated mast cells with poor outcomes^32–35^. However, none of these prior studies included within their analysis an evaluation of the gut microbiota or history of antibiotics exposure. Supporting our findings, McKee and colleagues demonstrated that antibiotic-induced perturbations of the gut microbiome significantly increased the density of mast cells in stromal-dense regions of luminal A and B tumors, resulting in increased tumor growth^10^. Although we did not observe differences in tumor growth with our model of dysbiosis, metastatic dissemination was significantly enhanced due to the accumulation of pro-fibrogenic mast cells. On a per-cell basis, we demonstrated that mast cells from mammary tissues of mice with established commensal dysbiosis were not only functionally distinct, they were sufficient to promote dissemination of HR^+^ tumors when implanted into mast cell-deficient mice. These findings highlight that antibiotic-induced perturbations to gut commensal homeostasis promote unfavorable outcomes for breast cancer through accumulation/activation of pro-fibrogenic mast cells.

Clinically, high levels of fibrosis are associated with increased invasiveness and poor outcomes in patients diagnosed with breast cancer^36^. However, until recently, the molecular mechanisms governing this process have remained unknown. Maller *et al*. recently uncovered a novel molecular pathway involving macrophage inflammation and collagen-mediated stiffening as a critical driver of tumor invasiveness and poor breast cancer outcomes^37^. Indeed, in the presence of a tumor, we observed increased macrophage numbers in adjacent tissues and tumors in mice with pre-established commensal dysbiosis^5^. Collagen levels in adjacent tissues and tumors were similarily increased^5^. However, macrophage numbers did not increase in pre-malignant mammary tissues in response to commensal dysbiosis alone. On the other hand, mast cell numbers were significantly increased in pre-malignant mammary tissues in response to dysbiosis and remained elevated in the presence of a tumor. Chemical inhibition of mast cells was sufficient to reduce collagen levels, suggesting that mast cells may also be involved in increasing tissue stiffness and matrix reorganization, at least in the context of changes to the microbiome.

Mast cell-mediated fibrosis is involved in multiple malignant and non-malignant pathologies; such as hepatocellular carcinoma^38^, intrahepatic cholangiocarcinoma^38^, and pulmonary fibrosis^39^. Commensal dysbiosis facilitates mast cell activation of mammary tissue fibroblasts, ultimately resulting in significantly increased tissue collagen I levels. Notably, increased protein levels of collagen I in the mammary tissues correlated with collagen levels in tumors of dysbiotic mice. These data mirror the findings of Arina *et al*. who demonstrated that tumor-associated SMA^+^ fibroblasts are recruited from the mammary tissue microenvironment^24^. Our findings indicate that mast cells are involved in this process through activation of tumor-associated fibroblasts in the adjacent mammary tissue, enhancing the reservoir of tumor-associated fibroblasts. Although we did not directly test whether fibroblast activation or collagen I deposition is the cause for increased tumor dissemination in response to commensal dysbiosis, we demonstrated that mast cells were sufficient for both activation of tissue fibroblasts and tumor dissemination. Based upon these data and the clinical relevance of fibrosis to invasive breast cancer, we hypothesize that mast cell-mediated fibrosis increases dysbiosis-induced tumor dissemination. Future work will interrogate the direct relevance of this pathway to dysbiosis-induced tumor dissemination.

Commensal dysbiosis increases platelet-derived growth factor (PDGF)-B^+^ expression in mast cells while also enhancing expression of the PDGF receptor PDGFR-*α* in fibroblasts. Mast cells are an important cellular source of PDGF, a well-known pro-fibrogenic signaling pathway. Indeed, mast cells sorted from mammary tissues of dysbiotic mice induced significantly higher levels of SMA and collagen I in PDGFR-a^+^ NIH-3T3 fibroblasts after *in vitro* co-culture, compared to mast cells from non-dysbiotic mice. Although PDGF signaling is associated with higher rates of breast cancer recurrence^40^, treatment with the PDGF receptor inhibitor imatinib has limited efficacy in patients with metastatic HR^+^ breast cancer^41–43^. Given the contribution of mast cells in the upregulation of PDGF signaling, imatinib therapy may benefit HR^+^ patients with a high abundance of PDGF^+^ mast cells in the adjacent mammary tissue or with commensal dysbiosis. Alternatively, mast cell stabilizers could potentially enhance the efficacy of imatinib for these patients.

Despite the understanding that bacterial and viral products can stimulate mast cells^44^, the mechanism by which intestinal microbiota influence mast cells in distal tissues remains largely unknown. Mechanistically, we demonstrate that blockade of CCL2 prior to tumor initiation was sufficient to reduce mast cell accumulation in the mammary tissue and diminish early dissemination of mammary tumor cells. These data uncover a potentially novel tumor-independent role for CCL2 in promoting mammary tumor progression and metastatic spread. Although CCL2 is a well-known driver of breast tumor growth^45^ and metastasis^17^, tumor-independent triggers of CCL2 have not been investigated. Supporting the idea that CCL2 contributes to tissue fibrosis in the absence of tumors and negative breast cancer outcomes when a tumor is present, Sun *et al*. demonstrated that overexpression of CCL2 in the pre-malignant mammary epithelium not only increased mammary tissue density, but also susceptibility to carcinogen-induced breast cancer^46^. Although the cell populations producing CCL2 in the mammary tissue in response to dysbiosis are undefined in our model, it is tempting to speculate that mast cells may be important for the fibrotic phenotype in the Sun *et al*. model. It would also be interesting to determine whether women with inflammatory breast disease also have an increased abundance of mast cells infiltrating into the tissues. Our analysis of transcriptomic data from normal breast tissues revealed gene enrichment of mast cell and cancer-associated fibroblast (CAF) subsets in patients who later developed breast cancer. These data suggest that, similar to CCL2^9^, increased abundance of mast cells is a negative correlate of breast cancer outcomes. Because the CCL2-CCR2 axis regulates the recruitment of mast cell progenitors (Mcps) during pulmonary inflammation^47^, future studies are aimed at defining whether the CCL2-CCR2 axis also regulates the recruitment of Mcps into mammary tissue, where they are matured into mast cells in response to early tumor growth.

While abundant evidence exists to elucidate tumor-intrinsic factors associated with breast cancer outcomes, emerging data reveals that genetic changes favoring tumor progression also occur in the adjacent mammary tissues^48^, even prior to breast cancer diagnosis^23^. These transcriptomic data suggest that tumor-independent and host-intrinsic factors negatively affect the mammary tissue environment to facilitate breast tumor metastasis. Gut commensal dysbiosis is sufficient to promote inflammation in adjacent mammary tissues and to facilitate dissemination of HR^+^ tumors in a murine model^5^. Recently, Terrisse *et al*. identified gut microbiota signatures in patients recently diagnosed with breast cancer that associated with tumor invasiveness and metastatic recurrence^6^. Microbes shown to promote negative outcomes in breast cancer were also found to be abundant in the gut microbiome of mice with commensal dysbiosis^5^, supporting that the gut microbiome contributes to negative breast cancer outcomes.

Breast cancer risk factors such as obesity, use of antibiotics, age, diet, and race, are all associated with altered composition of the commensal microbiome^49^. This highlights the necessity to define how an unbalanced commensal microbiome facilitates dissemination of breast tumor cells. There is a growing body of evidence indicating that the commensal microbiome affects multiple cancer outcomes and therapy responses^6,50, 51^. This study defines how an unbalanced and inflammatory microbiome promotes cellular and molecular changes within pre-malignant mammary tissue to facilitate early dissemination of HR^+^ tumors. Follow-up studies will further define how alterations to the commensal microbiome change the phenotype of mast cells residing in or recruited into the mammary tissue. Additionally, it will be important to determine whether commensal dysbiosis-driven phenotypic changes to mammary tissue mast cells result in other cellular and molecular changes in the mammary tissue that also favor tumor dissemination and growth, such as angiogenesis and immune suppression. Altogether, this study demonstrates that mast cells in mammary tissue are skewed into a pro-metastasis phenotype in response to commensal dysbiosis. While identifying dysbiosis in patients remains difficult, examining the phenotype of mast cells in the adjacent mammary tissues of patient samples could serve as an alternative approach to further interrogate how changes in the tissue environment affect breast cancer outcomes. Based on this study, targeting mast cells in patients with early-stage HR^+^ breast cancer could serve as an adjuvant to existing therapies aimed at reducing breast cancer recurrence.

## Materials and methods

### Mice

5-8-week-old female C57BL/6 mice were purchased from Charles River Laboratories. B6.Cg-*Kit^W-sh^*/HNihrJaeBsmJ (*Kit^w-sh/w-sh^*; Sash)^52–54^ mice were purchased from The Jackson Laboratory and maintained in-house at the University of Virginia. All animals were maintained in specific-pathogen-free barrier conditions. All experiments in this study were approved by the University of Virginia Institutional Animal Care and Use Committee.

### Tumor implantation

The poorly metastatic HR^+^ mouse mammary cancer cell line BRPKp110 has been described previously^2^. 5E5 BRPKp110 cells were injected orthotopically into the abdominal mammary fat pad. The highly metastatic mouse luciferase-expressing mammary cancer cell line PyMT was cloned from a metastatic tumor derived from the HR^+^ MMTV-PyMT model as previously described^20^. In experiments using this cell line, 1E5 PyMT cells were injected orthotopically into the abdominal mammary fat pad. Cell lines were authenticated by maintaining at less than four passages, monitoring of morphology, and testing for mycoplasma.

### Antibiotics administration

Commensal dysbiosis was initiated in mice by orally gavaging mice daily for 14 days with 100 μl of an antibiotic cocktail containing vancomycin (0.5 mg/ml, Gold Biotechnology), ampicillin (1 mg/ml) metronidazole (1 mg/ml), neomycin (1 mg/ml), and gentamicin (1 mg/ml) – all from Sigma Aldrich, as previously reported^5^. Vehicle-treated mice received 100 μl of water. Mice were then left untreated for 4 days to allow the establishment of commensal dysbiosis before tumor implantation.

Fecal microbiota transplantation (FMT) experiments were performed as previously described^5^. Briefly, dysbiosis was induced in donor mice as described above. Cecal contents from dysbiotic and non-dysbiotic vehicle-treated (water) animals were collected, homogenized, and frozen at −80°C in sterile 1:1 glycerol/PBS. Recipient mice were then orally gavaged with the antibiotic cocktail used for inducing dysbiosis for 7 days, followed by 3 or 4 consecutive days of oral gavage with 200 μl of either dysbiotic or non-dysbiotic cecal contents. After the final day of gavage, recipient mice were rested for 7 days before tumor initiation to allow for microbial engraftment.

### Treatment with mast cell stabilizer

Ketotifen (Sigma-Aldrich) solution was freshly prepared prior to each treatment by diluting in sterile water followed by passage through a 0.22 μM syringe filter. Mice were orally gavaged with 100 μl of ketotifen (10 mg/kg) or 100 μl of water daily beginning 1-day post tumor implantation, and continuing until day 5 post tumor.

### Flow cytometry

Mammary tissues, tumors, and tumor draining lymph nodes were collected, weighed, homogenized, and digested with collagenase type I (Gibco). All samples were stained with Zombie Aqua (BioLegend) and Fc receptor blocker anti-CD16/32 for differentiating live cells and inhibiting non-specific binding, respectively. Fibroblasts were identified by surface staining with anti-mouse CD45-PE-Cy7 (30-F11), CD31-PerCP-Cy5.5 (390), gp38-APC-Cy7 (8.1.1), and PDGFR-α-BV605 (APA5) and intracellular staining with antimouse smooth muscle actin (SMA)-AF647 (SPM32) and type I collagen-AF488 (1310-30). Mast cells and basophils were identified by surface staining with anti-mouse CD45-APC-Fire (30-F11), Lineage cocktail-Biotin (145-2C11; RB6-8C5; RA3-6B2; Ter-119; M1/70), CD49b-FITC (HMa2), cKit-BV650 (ACK2), FcεRIα-Biotin (MAR_1), CCR2-AF647 (SA203G11), CXCR4-PE/Dazzle (L276F12), CXCR3-PerCP-Cy5.5 (S18001A), Streptavidin-PerCP-Cy5.5, Streptavidin-PE-Cy7, and intracellular staining with antimouse PDGFB-AF488 (F-3). Myeloid cell infiltration was identified by surface staining with anti-mouse CD45-APC (30-F11), CD11b-PE-Cy7 (M1/70), Ly6C-APC-Cy7 (HK1.4), Ly6G-FITC (1A8), F4/80-PerCP-Cy5.5 (BM8), CD206-PE/Dazzle (C068C2), CD86-BV650 (GL-1). T cells, NK cells, and NKT cells were detected by surface staining with anti-mouse CD45-PE (30-F11), CD3-FITC (17A2), CD4-PE-Cy7 (GK1.5), CD8-APC-Cy7 (53-6.7), and NK1.1 (PK136).

Tumor cell dissemination was quantitated using single-cell suspensions from lungs and tumor-draining lymph nodes through surface staining with anti-mouse CD45 (30-F11, APC) and intracellular staining with anti-GFP (B-2, PE) or anti-luciferase (C-12, PE). Circulating tumor cells from the blood were isolated and quantitated as previously described^5, 55^. Briefly, equal volumes of EDTA-treated blood underwent red blood cell lysis, were plated into 6-well culture dishes, and incubated for 7 days. Cells were detached and stained as described for quantitation of disseminated tumor cells.

All antibodies were purchased from BioLegend with the exception of anti-SMA (Novus Biologicals), anti-type I collagen (SouthernBiotech), anti-GFP (Santa Cruz) and anti-luciferase (Santa Cruz). Counting beads (AccuCount, Spherotech) were used per manufacturer recommendations. Samples were run on an Attune NxT flow cytometer (Thermo Fisher Scientific) and analyzed with FlowJo software. Absolute numbers were quantitated per the manufacturer’s instructions.

### *In vitro* co-culture with mammary tissue mast cells and fibroblasts

CD45^+^ cKit^+^ FceRIa^+^ mast cells were sorted from mammary tissues from non-dysbiotic or dysbiotic C57BL/6 mice using a BD FACSAria Fusion flow sorter (Flow Cytometry Core Facility, University of Virginia). The sorted mast cells were then co-cultured with NIH-3T3 fibroblasts (Mast cell: NIH-3T3 fibroblast = 1:20) for 24 hours in serum-free DMEM at 37°C, 5% CO2. After 24 hours, the phenotype of NIH-3T3 fibroblasts was analyzed by surface staining of anti-mouse CD45-PE, PDGFR-α-BV605, and intracellular staining with SMA-AF647, and collagen I-AF488.

### Mast cell transfer experiment

Mast cells (CD45^+^ cKit^+^ FceRIa^+^ cells) were sorted from mammary tissues of non-dysbiotic or dysbiotic C57BL/6 mice as described above. Equal numbers of mammary tissue mast cells were reconstituted in 100 μl of PBS and orthotopically transferred to the mammary tissues (approximately ~300 mast cells/mammary tissue were injected for each experiment) of *Kit^w-sh/w-sh^* (Sash) mice. Eighteen hours after the transfer of sorted mast cells, 5E5 BRPKp110 tumor cells were injected into the same mammary fat pad of *Kit^w-sh/w-sh^* (Sash) recipient mice.

### Immunoblot

Mammary tissues and tumors were snap-frozen in liquid nitrogen and homogenized and sonicated in ice cold RIPA buffer (Thermo Fisher Scientific) supplemented with proteinase inhibitors (Thermo Fisher Scientific), per manufacturer instructions. Membranes were imaged using ChemiDoc Imaging System (Bio-Rad). Antibodies used were: anti-mast cell tryptase antibody (EPR9522, Abcam), anti-collagen I antibody (ab34710, Abcam), goat-anti rabbit IgG H&L (HRP) (Abcam), anti-PDGF-B (F-3, Santa Cruz), anti-GAPDH antibody (6C5, Santa Cruz), and mouse-IgGκ-HRP (Santa Cruz). The immunoblots were quantified using ImageJ software.

### Histology and microscopy

Tissues were fixed in neutral-buffered formalin, paraffin-embedded, and cut in 5 μm sections (Research Histology Core, University of Virginia). For detecting mast cells, slides were stained using 0.1% toluidine blue solution (pH 2.0-2.5). To investigate collagen deposition, slides were stained using PicroSirius Red (0.1% Direct Red 80 in saturated aqueous picric acid, Sigma-Aldrich). Collagen deposition was quantified using ImageJ software by calculating the stained area per section. The total numbers of mammary tissue mast cells were normalized to the area of mammary tissues. All the slides were imaged with a Slide Scanner (Leica).

### Transcriptome analysis of human samples

Normal breast tissues from women either prior to their BC diagnosis (susceptible normal, ^27^) or age-matched healthy women were obtained from the Susan G. Komen Tissue Bank at Indiana University Simon Comprehensive Cancer Center (Table S2). Subjects were recruited under a protocol approved by the Indiana University Institutional Review Board (IRB protocol number 1011003097 and 1607623663). Total RNA was extracted from fresh frozen breast tissues using 3 mm zirconium beads (Benchmark Scientific, cat.# D1032-30) and the AllPrep DNA/RNA/miRNA kit (Qiagen) as previously described^56^. Then, a cDNA library was prepared using the TruSeq Stranded Total RNA Kit (Illumina, San Diego, CA) and sequenced using Illumina HiSeq4000. Reads were adapter trimmed and quality filtered using Trimmomatic ver. 0.38^57^ setting the cutoff threshold for average base quality score at 20 over a window of 3 bases. Reads shorter than 20 bases post-trimming were excluded (parameters: LEADING:20 TRAILING:20 SLIDINGWINDOW:3:20 MINLEN:20). Cleaned reads mapped to Human genome reference sequence GRCh38.p12 with gencode v.28 annotation, using STAR version STAR_2.5.2b^58^. (parameters: --outFilterMultimapNmax 20 --seedSearchStartLmax 16 --twopassMode Basic). Gene expression was quantified by counting the number of reads mapped to the exonic regions of the genes using featureCounts tool ver. 2.0.0 of subread package (parameters: -s 2 -p -B -C).Differential Expression Analysis was performed using DESeq2 ver. 1.12.3. The *p*values for genes indicating the probabilities of no differential expression across the sample groups were corrected for multiple testing using Benjamini-Hochberg method.

### Availability of data and materials

The dataset including transcriptome profiling of susceptible breast samples and age-matched healthy controls is available in GEO (accession number GSE166044, https://www.ncbi.nlm.nih.gov/geo/query/acc.cgi?acc=GSE166044).

### Statistical analysis

Statistical analysis was performed using Prism software (GraphPad). Statistical difference between data sets was evaluated using two-tailed Mann-Whitney *U* test or unpaired t-test as indicated in the figure legends. Correlation analysis was performed using linear regression.

Gene Set Enrichment Analysis (GSEA) software version 4.1.0 (Broad Institute)^59^ was applied for the RNA-seq data from healthy and susceptible human donors in Fig.7. The gene signatures of mast cells^28^ and cancer-associated fibroblasts^29^ subsets are listed in Table S2. PDGF-B was also included in the analysis of mast cell signatures. The following options were selected in the software: numbers of permutations: 1000; chip platform: Human_ENSEMBL_Gene_ID_MSigDB.V7.4.chip; permutation type: gene_set. The results from GSEA are evaluated based on the enrichment scores (ES), which represents the gene set is overrepresented at the top or bottom of a ranked list of genes of interested. The normalized enrichment score (NES) represents the enrichment score that has been normalized to the size of gene sets. The false discovery rate (FDR q-value) represents the estimated probability of a false positive finding. The familywise-error rate (FWER p-value) stands for a probability that the NES represents a false positive finding.

P-values <0.05 were considered significant.

## Author contributions

**T. Feng:** Conceptualization, formal analysis, validation, investigation, visualization, methodology, writing-original draft. **F.N. Azar:** Investigation and editing. **C.B. Rosean:** Investigation and editing. **M.T. McGinty:** Investigation and editing. **A.M. Putelo:** Investigation and editing. **S. Koli:** Investigation. **N. Marino:** Investigation and editing. **R. German:** Investigation. **R. Podicheti:** Investigation and editing. **S.A. Dreger:** Investigation, analysis and editing. **W.J. Fowler:** Investigation, analysis and editing. **S. Greenfield:** Investigation. **S.D. Robinson:** Resources, analysis and editing. **M.R. Rutkowski:** Conceptualization, resources, analysis, visualization, supervision, funding acquisition, writing–original draft, writing–review and editing.

## Acknowledgments

This study was supported by a CCR grant from Susan G Komen CCR17483602 and 1R01CA253285 from the NCI, with aid from Grant #IRG 81-001-26 from the American Cancer Society. CBR was supported by T32AI007496-21, MTM was supported by T32CA009109-46. The authors would like to acknowledge the University of Virginia Research Histology, Biorepository and Tissue Research Facility, Flow Cytometry Core Facility, Center for Comparative Medicine, and the Carter Immunology Center. Additionally, we would like to thank Dr. Jose R. Conejo-Garcia for the BRPKp110 HR^+^ breast tumor cell line, Dr. Paula D. Bos for the PyMT cell line, Dr. Kimberly D. Kelly for the NIH-3T3 fibroblast cell line, and Dr. Sanja Arandjelovic for critical reading of the manuscript

