## Supplemental figures and legends for "Reciprocal interactions between the gut microbiome and mammary tissue mast cells promote metastatic dissemination of HR^+^ breast tumors"

Figure S1

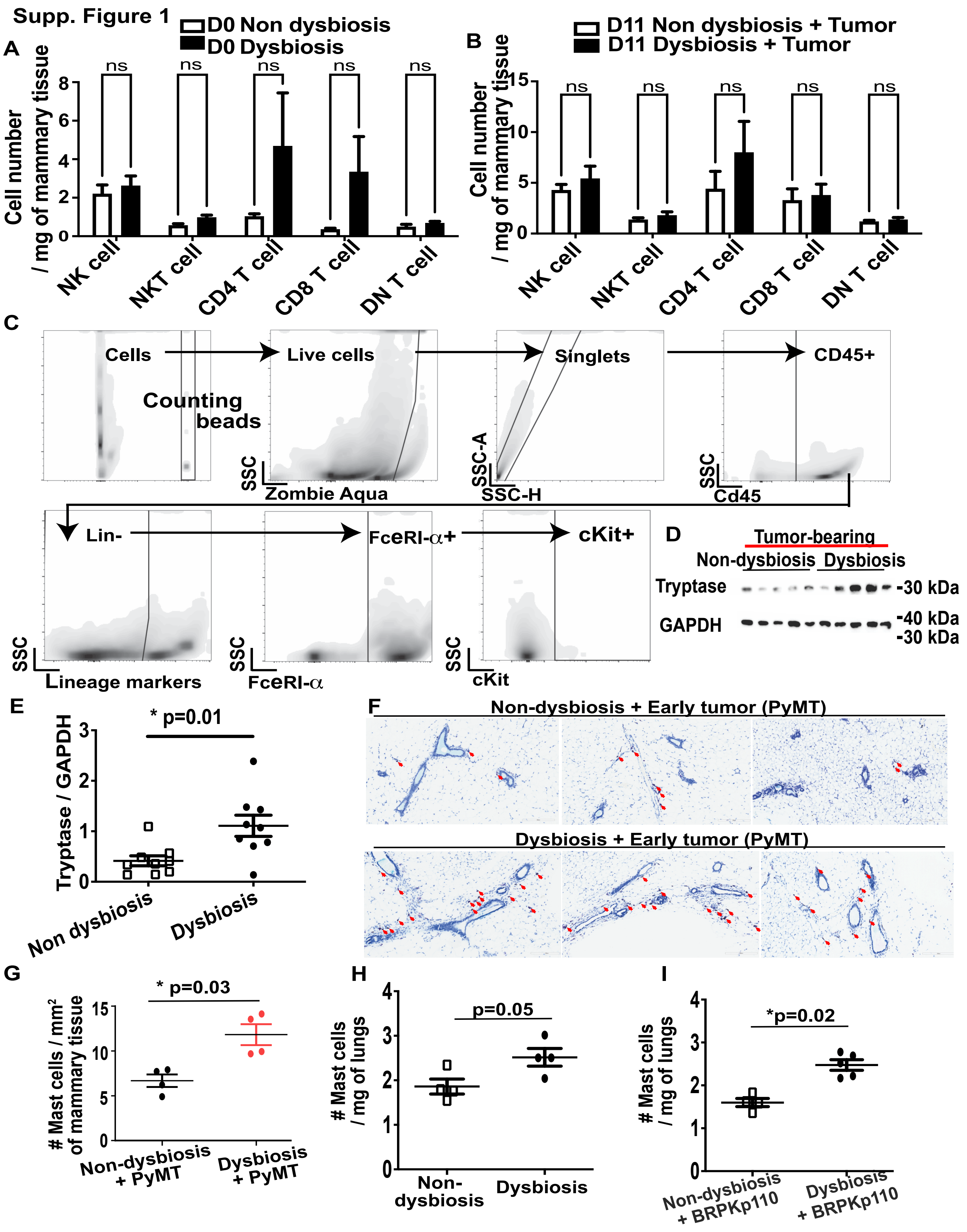
Supp. Figure 1
A
D0 Non dysbiosis
D0 Dysbiosis
Cell number / mg of mammary tissue
ns
NK cell
NKT cell
CD4 T cell
CD8 T cell
DN T cell
B
D11 Non dysbiosis + Tumor
D11 Dysbiosis + Tumor
C
Cells
Counting beads
Live cells
Singlets
CD45+
SSC
Zombie Aqua
SSC-A
SSC-H
Cd45
Lin-
FceRI-α+
cKit+
Lineage markers
FceRI-α
cKit
D
Tumor-bearing
Non-dysbiosis
Dysbiosis
Tryptase
GAPDH
-30 kDa
-40 kDa
-30 kDa
E
* p=0.01
Tryptase / GAPDH
Non dysbiosis
Dysbiosis
F
Non-dysbiosis + Early tumor (PyMT)
Dysbiosis + Early tumor (PyMT)
G
* p=0.03
### Mast cells / mm2 of mammary tissue
Non-dysbiosis + PyMT
Dysbiosis + PyMT
H
p=0.05
### Mast cells / mg of lungs
Non-dysbiosis
Dysbiosis
I
*p=0.02
Non-dysbiosis + BRPKp110
Dysbiosis + BRPKp110

**Mast cells accumulate in the mammary tissue in response to commensal dysbiosis and $HR^+$ mammary tumors.**

**A,** Mice were treated as in Figure 1A. On day 0, T cells and NK cells were enumerated in mammary tissues of dysbiotic and non-dysbiotic mice. **B,** Quantitation of T cells and NK cells from adjacent mammary tissues of tumor-bearing dysbiotic and non-dysbiotic mice 12 days after tumor initiation. For both A and B, all populations were gated on live, singlet, and CD45+. NK cells = $CD3^-$, $NK1.1^+$; NKT cells = $CD3^+$, $NK1.1^+$; CD4 T cells = $CD3^+$, $NK1.1^-$, $CD4^+$; CD8 T cells = $CD3^+$, $CD8a^+$, $NK1.1^-$; double negative (DN) T cells = $CD3^+$, $NK1.1^-$, $CD4^-$, $CD8a^-$. **C,** Gating strategy for identifying mast cells. Using flow cytometry, mast cells were identified as $CD45^+$ $Lin^-$ $FC\varepsilon RI\alpha^+$ $cKit^+$ cells. **D,** Immunoblot of the mast cell marker tryptase and GAPDH, as a loading control. Mammary tissues from dysbiotic or non-dysbiotic mice bearing day 12 BRPKp110 tumors were analyzed. **E,** Tryptase levels in adjacent mammary tissues were quantitated based on GAPDH levels using ImageJ. Graph represents values from two independent experiments. **F,** Representative images of adjacent mammary tissue isolated from mice bearing the aggressive $HR^+$ tumor PyMT 12 days post tumor initiation. Mast cells were identified from formalin fixed paraffin embedded mammary tissue whole mounts using toluidine blue. Red arrows indicate the mast cells. **G,** Quantification of mammary tissue mast cells. The total numbers of mast cells were enumerated from whole mammary tissue mounts obtained from mice bearing PyMT 12 days post tumor initiation. Mammary tissue mounts were sectioned and stained with Toluidine blue. Numbers of mast cells were normalized to the mammary tissue area. **H and I,** Numbers of mast cells in the lungs of non-tumor-bearing **(H)** and tumor-bearing mice **(I)** were evaluated by flow cytometry.

### Figure S2
#### Supp. Figure 2

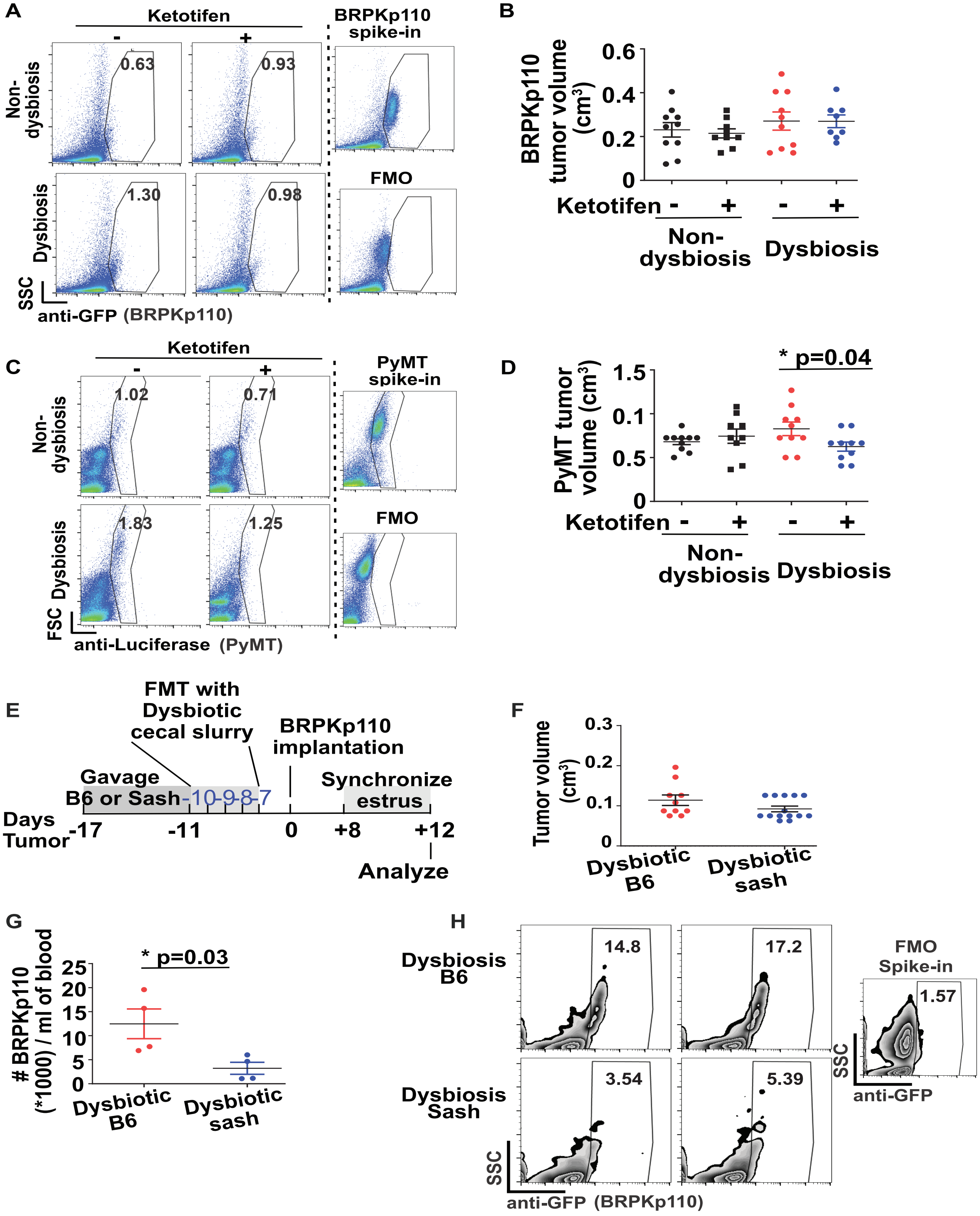

**Mast cells enhance dissemination of HR$^+$ tumor cells in response to a dysbiotic microflora.**

**A,** Representative plots demonstrating GFP$^+$ BRPKp110 cells in the lungs of mice 22 days post tumor initiation treated with or without Ketotifen. **B,** Final tumor volumes of BRPKp110 tumor-bearing mice treated with or without Ketotifen 22 days post tumor-initiation (see experimental layout Figure 2A). **C,** Representative plots demonstrating luciferase$^+$ PyMT cells in the lungs of mice 22 days post tumor initiation treated with or without Ketotifen. **D,** Final tumor volumes of PyMT tumor-bearing mice treated with or without Ketotifen 22 days post tumor-initiation (see experimental layout Figure 2A). For **A** and **C**, the gating strategy for tumor cells was determined based on a lung sample spiked with tumor cells and the fluorescence minus one (FMO) controls. Numbers represent percent of gated tumor cells of total live cells. **E,** Experimental design of F to G. Wild-type C57BL/6 (B6) mice and $Kit^{w\text{-}sh/w\text{-}sh}$ (Sash) mice were gavaged with a cocktail of broad-spectrum antibiotics for 7 days followed by fecal transfer of dysbiotic cecal slurries (from wild-type C57BL/6 mice) for 4 consecutive days. The microbiome was allowed to establish for 7 days, followed by initiation of tumors using BRPKP110 tumor cells. **F to H,** 12 days post tumor implantation, tumor volumes were evaluated **(F)**. Disseminated tumor cells in the peripheral blood were evaluated by flow cytometry **(G and H)**. **G,** Numbers of GFP$^+$ tumor cells in peripheral blood. **H,** Representative density flow cytometry plots showing GFP$^+$ BRPKP110 tumor cells in the peripheral blood. FMO spike-in represents a gating control of blood or lymph nodes spiked with BRPKP110 tumor cells, minus the anti-GFP antibody. Numbers represent the percentage of gated GFP$^+$ tumor cells of CD45 negative cells. Each symbol represents an individual mouse, and statistical significance was determined by two-tailed Mann-Whitney *U* test.

**Figure S3**

**Supp. Figure 3**

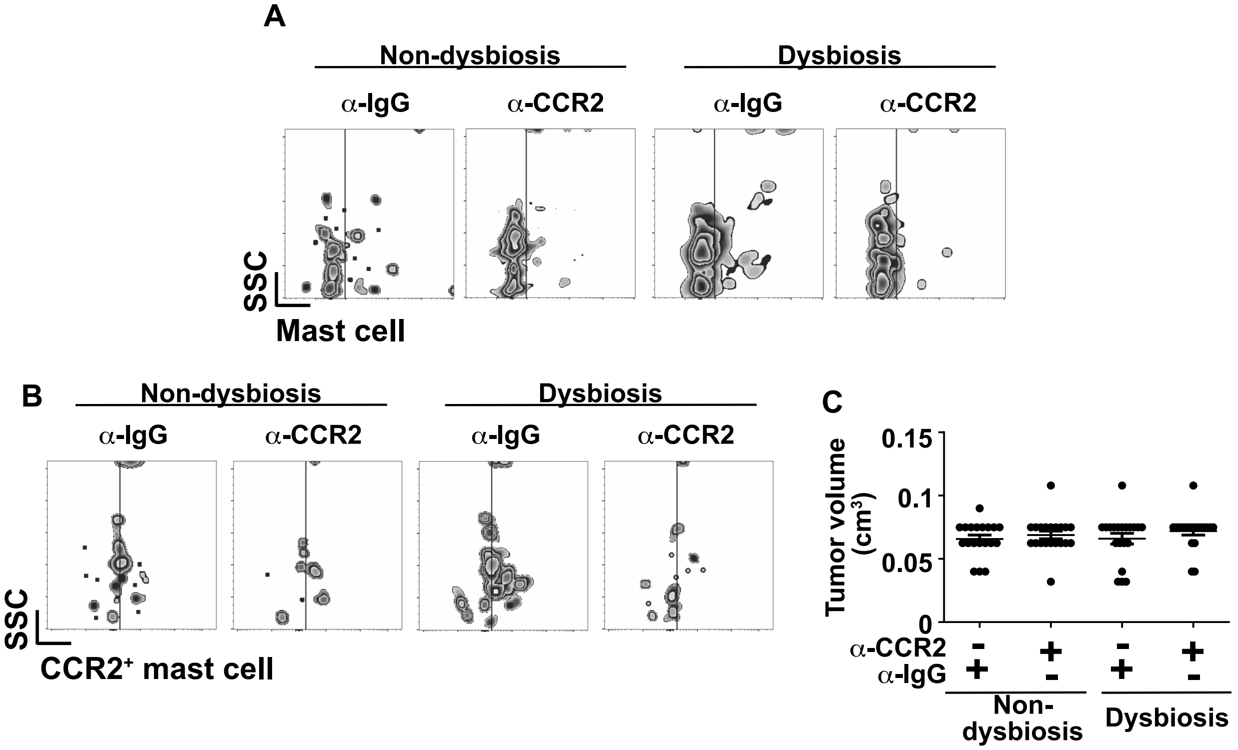


**Commensal dysbiosis-induced CCL2 promotes mast cell accumulation into the mammary tissue.**

**A and B,** Mice were intraperitoneally injected with 4 doses of anti-CCL2 or an isotype-matched control IgG prior to tumor initiation on day -8, day -6, day -4 and day -2. Tumors were initiated on day 0. Mammary tissues were analyzed on day 12, similar to experimental schematic in Figure 3B. Representative plots demonstrating total mast cells **(A)** and $CCR2^+$ mast cells **(B)** in the mammary tissues of mice in Figure 3C and Figure 3D. **C,** Tumor volumes of anti-CCL2 antibody- or IgG-isotype treated mice 12 days post tumor implantation**.** Each symbol represents an individual tumor, and statistical significance was determined by two-tailed Mann-Whitney *U* test.

Figure S4

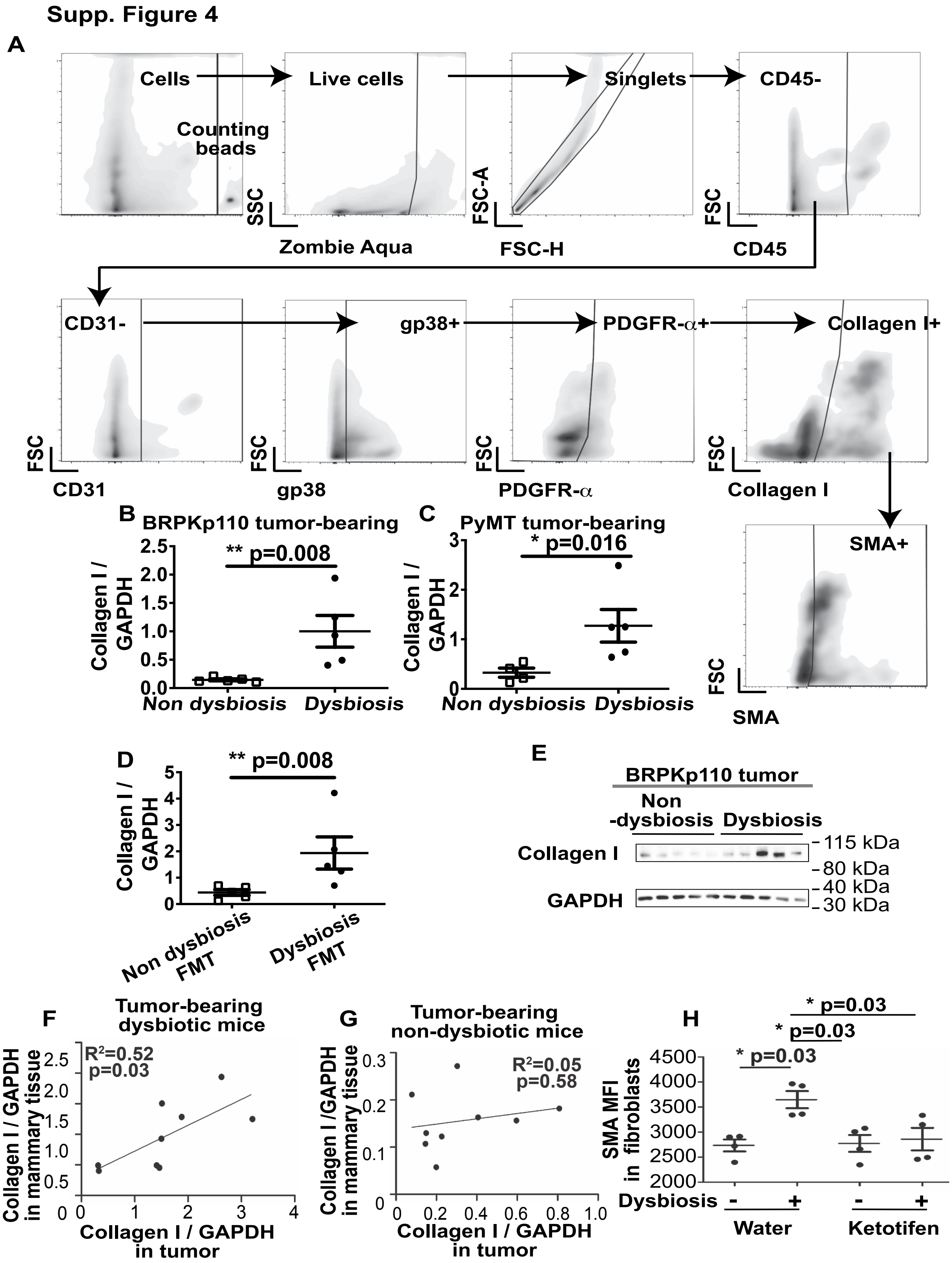
Supp. Figure 4
A
Cells
Counting beads
Live cells
Singlets
CD45-
SSC
Zombie Aqua
FSC-A
FSC-H
FSC
CD45
CD31-
gp38+
PDGFR-α+
Collagen I+
FSC
CD31
gp38
PDGFR-α
Collagen I
SMA+
SMA
B
BRPKp110 tumor-bearing
** p=0.008
Collagen I / GAPDH
Non dysbiosis
Dysbiosis
C
PyMT tumor-bearing
* p=0.016
Collagen I / GAPDH
Non dysbiosis
Dysbiosis
D
** p=0.008
Collagen I / GAPDH
Non dysbiosis FMT
Dysbiosis FMT
E
BRPKp110 tumor
Non -dysbiosis
Dysbiosis
Collagen I
GAPDH
115 kDa
80 kDa
40 kDa
30 kDa
F
Tumor-bearing dysbiotic mice
R²=0.52
p=0.03
Collagen I / GAPDH in mammary tissue
Collagen I / GAPDH in tumor
G
Tumor-bearing non-dysbiotic mice
R²=0.05
p=0.58
Collagen I /GAPDH in mammary tissue
Collagen I / GAPDH in tumor
H
* p=0.03
* p=0.03
* p=0.03
SMA MFI in fibroblasts
Dysbiosis
Water
Ketotifen

**Protein levels of collagen I in the mammary tissue is positively correlated with collagen I protein levels in the tumors of mice with commensal dysbiosis, but not in mice without dysbiosis.**

**A,** Gating strategy for identifying collagen $I^+$ $SMA^+$ fibroblasts. **B and C,** Collagen I levels in the mammary tissue were evaluated by immunoblot 12 days post BRPKp110 tumor implantation **(B)** or PyMT tumor implantation **(C)** and quantified based on GAPDH levels using ImageJ. **D,** Mice received a fecal transplant of non-dysbiotic or dysbiotic cecal slurries prior to tumor initiation, as in Figure 1G. Levels of collagen I protein in the mammary tissue were quantitated based on GAPDH protein levels using ImageJ 12 days post tumor implantation with BRPKp110. **E,** Immunoblotting of collagen I and GAPDH in tumors from non-dysbiotic and dysbiotic mice. **F and G,** Linear correlation of collagen I protein levels in the adjacent mammary tissues and tumors from dysbiotic mice **(F)** and non-dysbiotic mice **(G)**. **H,** Mean fluorescence intensity (MFI) of SMA in mammary tissue fibroblasts ($CD45^-$ $CD31^-$ $gp38^+$ PDGFR-$\alpha^+$ cells) from tumor-bearing non-dysbiotic or dysbiotic mice after treatment with the mast cell stabilizer ketotifen or water, similar to Fig. 4H. Each symbol represents an individual mouse, and statistical significance was determined by linear correlation (C and D) or two-tailed Mann-Whitney *U* test (E).

#### Figure S5

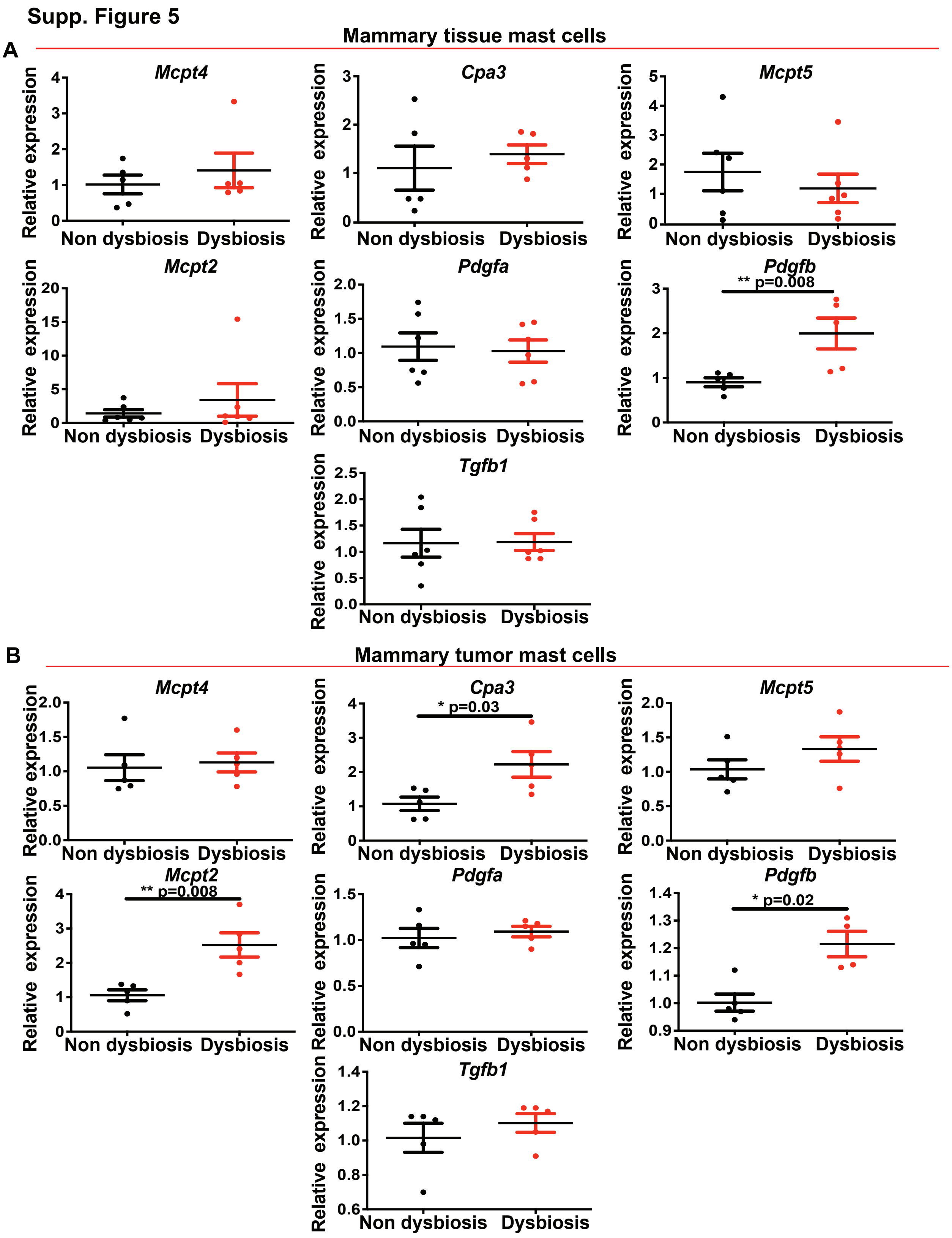

Supp. Figure 5
A
Mammary tissue mast cells
Mcpt4
Cpa3
Mcpt5
Mcpt2
Pdgfa
Pdgfb
** p=0.008
Tgfb1
Relative expression
Non dysbiosis
Dysbiosis
B
Mammary tumor mast cells
Mcpt4
Cpa3
* p=0.03
Mcpt5
Mcpt2
** p=0.008
Pdgfa
Pdgfb
* p=0.02
Tgfb1

**Mast cells from mammary tissues of dysbiotic tumor-bearing mice are phenotypically distinct from mast cells in mammary tissues of non-dysbiotic mice.**

Mast cells were sorted from mammary tissues of dysbiotic or non dysbiotic mice as described above. Total RNA was extracted from using a RNeasy Micro Kit (Qiagen) followed by reverse transcription using High-Capacity cDNA Reverse Transcription Kits (Thermo Fisher). Primers are listed in table S1. For amplicon detection, we used the Power SYBR® Green PCR Master Mix (Thermo Fisher) as described by the manufacturer. RT-PCR was performed in a QuantStudio 6 Real-Time PCR Systems (Thermo Fisher) as follows: initial denaturation at 95°C for 10 mins; amplification for 35 cycles of denaturation (95°C for 1 mins), annealing (55°C for 2 mins), and extension (72°C for 3 mins). Specificity of the amplicon was determined by melting curve. The relative levels of mRNA were determined by comparative CT method and normalized by housekeeping genes GAPDH and β-actin RNA.

The following primers were used: *Pdgfa*: Forward primer 5'- GTGCGACCTCCAACCTGA-3' and reverse primer 5'- GGCTCATCTCACCTCACATCT; *Pdgfb*: Forward primer 5'-CGGCCTGTGACTAGAAGTCC-3' and reverse primer 5'- GAGCTTGAGGCGTCTTGG-3'; *Tgf-β1* forward primer 5'- TGGAGCAACATGTGGAACTC-3' and reverse primer 5'-CAGCAGCCAATTACCAAG-3'; *Cpa3* forward primer 5'- ATCGCAGGCACGCACAGTTAT-3' and reverse primer 5'- AACCCAGTCTAAGGAAGAGCC-3'; *Mcpt4* forward primer 5'-ACCACTGAGAGAGGGTTCACAGC-3' and reverse primer 5'-GAAGACTCTGATGCACGCAGG-3'; *Mcpt5* forward primer 5'-CTGAGAACTACCTGTCGGCCTGC-3' and reverse primer 5'-TCCAGTTCCAGATTTCCTCACGG-3'; *Mcpt2* forward primer 5'-CCACTAAGAACGGTTCGAAGGAG-3' and revere primer 5'-GCTGGGATGAACTCAGAGGTACC-3'; *β-actin* forward primer 5'-GTGGGCCGCTCTAGGCACCAA-3' and reverse primer 5'-CTCTTTGATGTCACGCACGATTTC-3'; *Gapdh* forward primer 5'-CATCACTGCCACCCAGAAGACTG-3' and reverse primer 5'-ATGCCAGTGAGCTTCCCGTTCAG-3'

Commensal dysbiosis leads to the accumulation of mast cells into mammary tissues of tumor-bearing mice with increased expression levels of the pro-fibrogenic mediator platelet derived growth factor subunit B (*Pdgfb*). Mast cells were sorted from adjacent mammary tissues **(A)** or BRPKp110 mammary tumors **(B)** of dysbiotic or non-dysbiotic tumor-bearing mice during early (day 6) tumor progression. RNA was extracted from sorted cells and relative expression of each gene was measured using semi-quantitative PCR. Relative gene expression levels were calculated based upon CT value for each gene/sample and normalized to the averaged CT values of GAPDH and β-actin. Each symbol represents an experimental replicate, and statistical significance was determined by two-tailed Mann-Whitney *U* test.

#### Figure S6

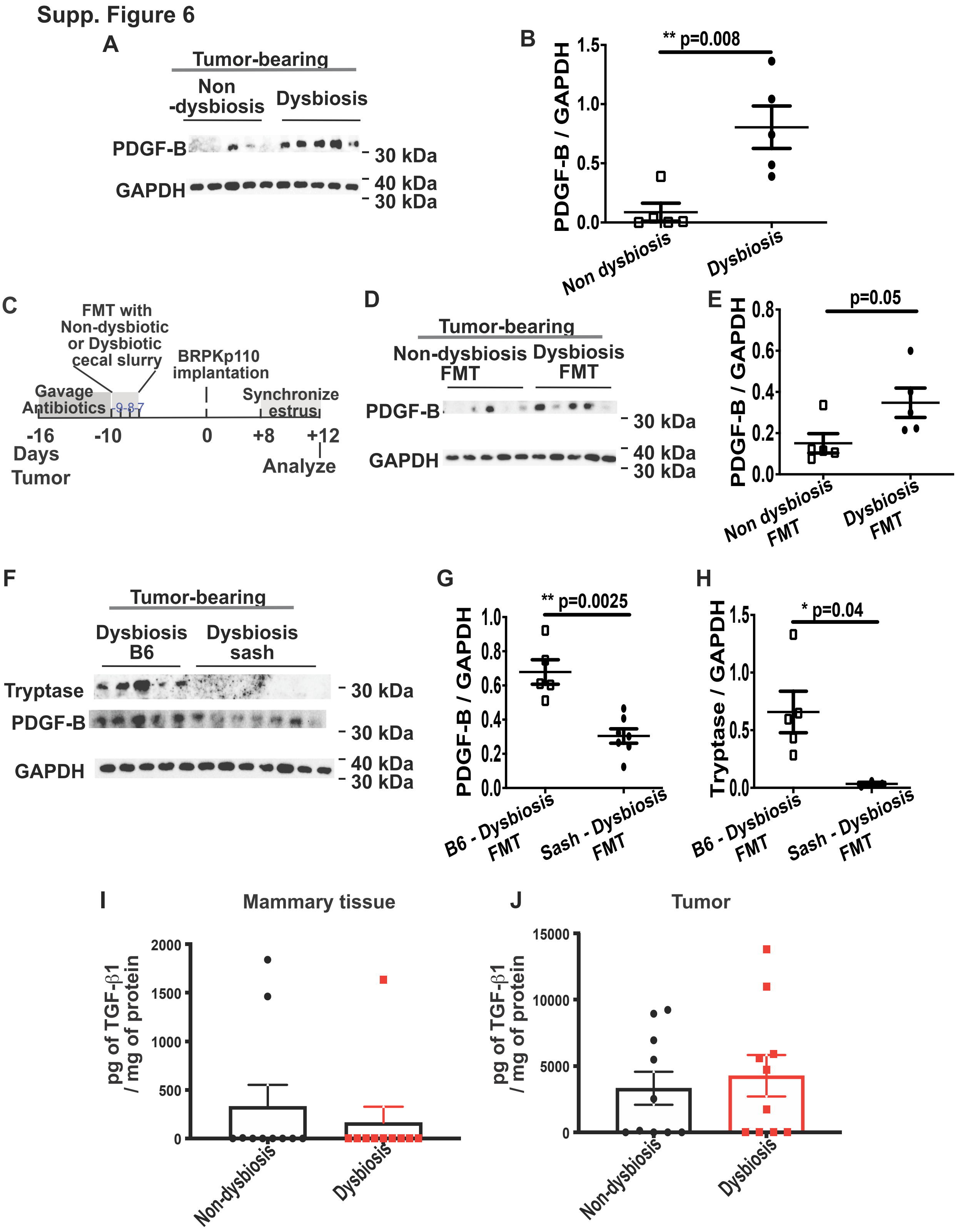
Supp. Figure 6
A
Tumor-bearing
Non -dysbiosis
Dysbiosis
PDGF-B
GAPDH
30 kDa
40 kDa
30 kDa
B
** p=0.008
PDGF-B / GAPDH
Non dysbiosis
Dysbiosis
C
FMT with Non-dysbiotic or Dysbiotic cecal slurry
BRPKp110 implantation
Gavage Antibiotics
Synchronize estrus
-16
-10
0
+8
+12
Days Tumor
Analyze
D
Tumor-bearing
Non-dysbiosis FMT
Dysbiosis FMT
PDGF-B
GAPDH
E
p=0.05
PDGF-B / GAPDH
Non dysbiosis FMT
Dysbiosis FMT
F
Tumor-bearing
Dysbiosis B6
Dysbiosis sash
Tryptase
PDGF-B
GAPDH
G
** p=0.0025
PDGF-B / GAPDH
B6 - Dysbiosis FMT
Sash - Dysbiosis FMT
H
* p=0.04
Tryptase / GAPDH
B6 - Dysbiosis FMT
Sash - Dysbiosis FMT
I
Mammary tissue
pg of TGF-β1 / mg of protein
Non-dysbiosis
Dysbiosis
J
Tumor
pg of TGF-β1 / mg of protein
Non-dysbiosis
Dysbiosis

**Commensal dysbiosis increases protein levels of profibrogenic mediator PDGF-B in the mammary tissues and on mammary tissue-associated mast cells of tumor-bearing mice.**

**A**, Immunoblot of PDGF-B from mammary tissue lysates of BRPKp110 tumor-bearing mice with or without commensal dysbiosis. **B,** PDGF-B levels in the mammary tissue were evaluated by immunoblot 12 days post BRPKp110 tumor implantation and quantified based on GAPDH levels using ImageJ. **C,** Experimental schematic for D and E. Mice received a fecal transplant of non-dysbiotic or dysbiotic cecal slurries prior to tumor initiation, as in Figure 1G. **D,** Immunoblot of PDGF-B or GAPDH as a protein loading control in mice receiving a fecal transfer (FMT) with normal or dysbiotic fecal slurry. **E**, Quantitation of PDGF-B levels in the mammary tissue based on GAPDH levels using ImageJ. **F,** Tumor-bearing dysbiotic C57BL/6 mice (B6) and mast cell-deficient $\textit{Kit}^{w\text{-}sh/w\text{-}sh}$ mice (Sash) were reconstituted with a dysbiotic cecal flora as demonstrated in B. Mammary tissues were analyzed 12 days post tumor initiation. Protein levels of tryptase and PDGF-B in the mammary tissues were determined by western blot. GAPDH served as a protein loading control. **G,** Quantitation of mammary tissue PDGF-B levels from the blot in F, based on GAPDH levels using ImageJ. **H,** Quantitation of mammary tissue Tryptase levels from the blot in F based on GAPDH levels using ImageJ. **I and J,** TGF$\beta$ protein levels from paired mammary glands **(I)** or tumors **(J)** of non-dysbiotic and dysbiotic mice 12 days post tumor initiation. Each symbol represents an individual mouse, and statistical significance was determined by two-tailed Mann-Whitney *U* test.
